# Substitution of nucleotide-sugar by trehalose-dependent glycogen synthesis pathways in Chlamydiales underlines an unusual requirement for storage polysaccharides within obligate intracellular bacteria

**DOI:** 10.1101/2020.06.02.131169

**Authors:** Matthieu Colpaert, Derifa Kadouche, Mathieu Ducatez, Trestan Pillonel, Carole Kebbi-Beghdadi, Ugo Cenci, Binquan Huang, Malika Chabi, Emmanuel Maes, Bernadette Coddeville, Loïc Couderc, Hélène Touzet, Fabrice Bray, Catherine Tirtiaux, Steven Ball, Gilbert Greub, Colleoni Christophe

**Affiliations:** University of Lille, CNRS, UMR8576-UGSF-Unité de Glycobiologie Structurale et Fonctionnelle, F-59000 Lille, France; Institute of Microbiology, University of Lausanne and University Hospital Center, Lausanne, Switzerland; University of Lille, CNRS, Inserm, CHU Lille, Institut Pasteur de Lille, US 41 - UMS 2014 - PLBS, F-59000 Lille, France; University of Lille, CNRS, Centrale Lille, UMR 9189 - CRIStAL - Centre de Recherche en Informatique Signal et Automatique de Lille, F-59000 Lille, France; University of Lille, CNRS, USR 3290—MSAP—Miniaturisation pour la Synthèse, l’Analyse et la Protéomique, F-59000 Lille, France

## Abstract

All obligate intracellular pathogens or symbionts of eukaryotes lack glycogen metabolism. Most members of the *Chlamydiales* order are exceptions to this rule as they contain the classical GlgA-GlgC-dependent pathway of glycogen metabolism that relies on the ADP-Glucose substrate. We surveyed the diversity of Chlamydiales and found glycogen metabolism to be universally present with the important exception of *Criblamydiaceae* and *Waddliaceae* families that had been previously reported to lack an active pathway. However, we now find elements of the more recently described GlgE maltose-1-P-dependent pathway in several protist-infecting Chlamydiales. In the case of *Waddliaceae* and *Criblamydiaceae*, the substitution of the classical pathway by this recently proposed GlgE pathway was essentially complete as evidenced by the loss of both GlgA and GlgC. Biochemical analysis of recombinant proteins expressed from *Waddlia chondrophila* and *Estrella lausannensis* established that both enzymes do polymerize glycogen from trehalose through the production of maltose-1-P by TreS-Mak and its incorporation into glycogen’s outer chains by GlgE. Unlike *Mycobacteriaceae* where GlgE-dependent polymerization is produced from both bacterial ADP-Glc and trehalose, glycogen synthesis seems to be entirely dependent on host supplied UDP-Glc and Glucose-6-P or on host supplied trehalose and maltooligosaccharides. These results are discussed in the light of a possible effector nature of these enzymes, of the chlamydial host specificity and of a possible function of glycogen in extracellular survival and infectivity of the chlamydial elementary bodies. They underline that contrarily to all other obligate intracellular bacteria, glycogen metabolism is indeed central to chlamydial replication and maintenance.

## Introduction

*Chlamydiae* forms with Planctomycetes and Verrucomicrobia phyla a very ancient monophyletic group of bacteria known as PVC, which has been recently enriched with additional phyla [1]. The *Chlamydiales* order groups the members of the *Chlamydiaceae* family that includes etiological agents of humans and animals infectious diseases and at least eight additional families commonly named “*chlamydia*-related bacteria or “environmental” chlamydia [2, 3].

The hallmark of Chlamydiales consists in an obligate intracellular lifestyle due to a significant genome reduction and biphasic development, which includes two major morphological and physiological distinct stages: the elementary body (EB), a non-dividing and infectious form adapted to extracellular survival and the reticulate body (RB), a replicating form located within a membrane surrounded inclusion (for review [4]). Following entry into a susceptible cell the EBs differentiate into RBs within the inclusion. During the intracellular stage, RBs secrete many effector proteins through the type III secretion system and express a wide range of transporters in order to manipulate host metabolism and uptake all the metabolites required for its replication. At the end of the infection cycle, RBs differentiate back into EBs before they are released into the environment [5, 6].

Glycogen metabolism loss appears to be a universal feature of the reductive genome evolution experienced by most if not all obligate intracellular bacterial pathogens or symbionts including *Anaplasma* spp., *Ehrlichia* spp., *Wolbachia* spp., *Rickettsia* spp. (alpha-proteobacteria), *Buchnera* sp. *Coxiella* sp. (gamma-proteobacteria), or *Mycobacterium leprae* (Terrabacteria) [7, 8]. Despite the more advanced genome reduction experienced by the animal-specific *Chlamydiaceae* family (0.9 Mpb) in comparison to other protist-infecting Chlamydiales (2 to 2.5 Mpb), the glycogen metabolism pathway appears surprisingly preserved [7]. This includes the three-enzymatic activities required for glycogen biosynthesis: GlgC, GlgA and GlgB. ADP-glucose pyrophosphorylase (GlgC) activity that controls the synthesis and level of nucleotide-sugar, ADP-glucose, dedicated solely to glycogen biosynthesis. Glycogen synthase (GlgA) belongs to the Glycosyl Transferase 5 family (GT5: CaZy classification) which polymerizes nucleotide-sugar into linear α-1,4 glucan.

GlgA activity has a dual function consisting of a primer-independent glucan synthesis and glucan elongation at the non-reducing end of preexisting polymers [9]. When the primer reaches a sufficient degree of polymerization (DP>15) to fit the catalytic site of the glycogen branching enzyme (GlgB), α-1,6 branches are introduced resulting in the appearance of two non-reducing polymer ends that may be further elongated by GlgA. The repetition of this process results in an exponential increase in the number of non-reducing ends leading to a particle with a 32-40 nm diameter [10].

Until recently, *Waddlia chondrophila* (family *Waddliaceae*) as well as all members of *Criblamydiaceae* could be considered as important exceptions to the universal requirement of *Chlamydiales* for glycogen synthesis. Indeed, genome analysis indicated that *ad minima* the *glgC* gene was absent from all these bacteria [11–13] and that the function of GlgA was possibly also impaired. Consequently, based on the absence of glycogen reported for all knockout *glgC* mutants in bacteria and plants it was believed that *W. chondrophila* was defective in glycogen synthesis [14, 15]. Using transmission electron microscope analysis, we are now reporting numerous glycogen particles within the cytosol of *W. chondrophila* and *Estrella lausannensis* (family *Criblamydiaceae*) EBs, suggesting either another gene encodes a phylogenetically distant protein that overlaps the GlgC activity or an alternative glycogen pathway takes place in these Chlamydiales.

The recent characterization of an alternative glycogen pathway, the so-called GlgE-pathway, in *Mycobacterium tuberculosis* and streptomycetes prompted us to probe chlamydial genomes with homolog genes involved in this pathway [16, 17]. At variance with the nucleotide-sugar based GlgC-pathway, the GlgE-pathway consists of the polymerization of α-1,4 glucan chains from maltose 1-P. In Mycobacteria, the latter is produced either from the condensation of glucose-1-P and ADP-glucose catalyzed by a glycosyl transferase called GlgM or from the interconversion of trehalose (α-α−1,1 linked D-glucose) to maltose followed by the phosphorylation of maltose, which are catalyzed by trehalose synthase (TreS) and maltose kinase (Mak) activities, respectively [16]. At the exception of Actinobacteria (i.e mycobacteria and Streptomycetes), TreS is usually fused with a maltokinase (Mak) that phosphorylates maltose into maltose-1-phosphate [18]. Subsequently, maltosyl-1-phosphate transferase (GlgE) mediates the formation of α-1,4-linked polymers by transferring the maltosyl moiety onto the non-reducing end of a growing α-1,4-glucan chain. As in the GlgC-pathway, branching enzyme (GlgB) introduces α-1,6 linkages to give rise to a highly branched α-glucan. The GlgC-pathway is found in approximately one third of the sequenced bacteria and is by far the most widespread and best studied; the GlgE pathway has been identified in 14% of the genomes of α-, β-γ-proteobacteria while 4% of bacterial genomes possess both GlgC- and GlgE-pathways [18, 19].

In order to shed light on the metabolism of storage polysaccharide in Chlamydiales, we analyzed 220 genomes (including some genomes assembled from metagenomic data) from 47 different chlamydial species that represent the bulk of currently known chlamydial diversity. A complete GlgE-pathway was identified in five chlamydial species distributed in *Criblamydiaceae*, *Waddliaceae* and *Parachlamydiaceae* families. In this work, we demonstrated that the GlgC-pathway is impaired in C*riblamydiaceae* and *Waddliaceae*. The complete biochemical characterization of the GlgE-pathway in *Estrella lausannensis* (family *Criblamydiaceae*) and *Waddlia chondrophila* (family *Waddliaceae*) isolated respectively from water in Spain [20, 21] and from the tissue of an abortive bovine fetus [22, 23] is reported. Thus, despite the intensive reductive genome evolution experienced by these intracellular bacteria our work shows that glycogen biosynthesis is maintained in all Chlamydiales and suggests a hitherto understudied function of storage polysaccharides and oligosaccharides in the lifecycle of all *Chlamydiales*.

## Materials and methods

### Comparative genomic analysis of glycogen metabolic pathways

In order to gain insight into Chlamydiae’s glycogen metabolism, homologs of proteins part of the glycogen pathway of *E.coli* and of *M. tuberculosis* were searched with BLASTp in 220 genomes and metagenome-derived genomes from 47 different chlamydial species available on the ChlamDB database (https://chlamdb.ch/, https://academic.oup.com/nar/article/48/D1/D526/5609527)[24]. The completeness of metagenome-derived and draft genomes was estimated with checkM based on the identification of 104 nearly universal bacterial marker genes [3]. The species phylogeny has also been retrieved from ChlamDB website.

### Microscopy analysis

Fresh cultures of *Acanthamoeba castellanii* grown in 10 mL YPG (Yeast extract, peptone, glucose) were infected with one-week-old 5 µm-filtered suspension of *E. lausannensis* or *W. chondrophila* (10^5^ cells.mL^-1^), as previously reported [25].Samples of time course infection experiments were harvested at 0, 7, 16 and 24 hours post-infection by centrifuging the infected *A. castellanii* cultures at 116 g. Pellets were then fixed with 1 mL of 3 % glutaraldehyde for four hours at 4°C and prepared as described previously [26].

### Glycogen synthase and glycogen branching enzyme activities

Fused *glgAglgB* genes of *E. lausannensis* and *W. chondrophila* were amplified using primer couples harboring attB sites as described in the **S1 table**. PCR products were then cloned in the pET15b (Novagen) plasmid modified compatible with Gateway^TM^ cloning strategy. The expression of his-tagged recombinant protein GlgA-GlgB was performed in the derivative BW25113 strain impaired in the endogenous glycogen synthase activity (Δ*glgA*). Glycogen synthases assay and zymogram analysis have been conducted as described previously [27]. The nucleotide-sugar specificity of glycogen synthase was carried out by following the incorporation of ^14^C-Glc of radiolabelled ADP-^14^C-[U]-glc or UDP-^14^C-[U]-glc into glycogen particles during one hour at 30°C.

### Phylogeny analysis

Homologous sequences of TreS-Mak and GlgE were carried out by BLAST against the nr database from NCBI with respectively WP_098038072.1 and WP_098038073.1 sequences of *Estrella lausannensis*. We retrieved the top 2000 homologs with an E-value cut off ≤ 10 ^-5^ and aligned them using MAFFT [28] with the fast alignment settings. Block selection was then performed using BMGE [29] with a block size of 4 and the BLOSUM30 similarity matrix. Preliminary trees were generated using Fasttree [30] and ‘dereplication’ was applied to robustly supported monophyletic clades using TreeTrimmer [31] in order to reduce sequence redundancy. For each protein, the final set of sequences was selected manually. Proteins were re-aligned with MUSCLE [32] block selection was carried out using BMGE with a block size of four and the matrix BLOSUM30, and trees were generated using Phylobayes [33] under the catfix C20 + Poisson model with the two chains stopped when convergence was reached (maxdiff<0.1) after at least 500 cycles, discarding 100 burn-in trees. Bootstrap support values were estimated from 100 replicates using IQ-TREE [34] under the LG4X model and mapped onto the Bayesian tree.

### GlgE and TreS-Mak expressions

*glgE* and *treS-mak* genes were amplified from the genomic DNA of *E. lausannensis* and *W. chondrophila* by the primers F_glgE_EL/R_glgE_EL, F_glgE_WC/R_glgE_WC and F_treS-mak_EL/R_treS-mak_EL (**S1 table**). The PCR products were cloned in the expression vector pET15b (Novagene) or VCC1 (P15A replicon). The resulting plasmids pET-GlgE-WC, pET-GlgE-EL, VCC1-treS-mak-EL were transferred to *E. coli* Rosetta™ (DE3; pRARE) or BL21-AI™. The expression of his-tagged proteins was induced in Lysogeny Broth (LB) or Terrific Broth (TB) by the addition of 0.5 mM isopropyl β-D-1-thiogalactopyranoside (IPTG) or 1 mM IPTG/0.2 % L-arabinose at the mid-logarithmic phase growth (A600=0.5), or by using auto inducible medium as described in Fox and Blommel [35]. After 18h at 30°C, cells were harvested at 4000g at 4°C during 15 minutes. Cell pellets were stored at −80°C until purification step on Ni ^2+^ affinity column.

### Protein purification

Cell pellets from 100 mL of culture medium were resuspended in 1.5 mL of cold buffer (25 mM Tris-acetate, pH 7.5). After sonication (three times 30 s), proteins were purified on Ni ^2+^ affinity column (Roth) equilibrated with washing buffer (300 mM NaCl, 50 mM sodium acetate and 60 mM imidazole, pH 7) and eluted with a similar buffer containing 250 mM imidazole. Purification steps were followed by SDS-PAGE and purified enzymes quantified by Bradford method (Bio-Rad).

### Evidence of GlgE-like activity by thin layer chromatography

Maltosyl transferase activities of GlgE proteins of *E. lausannensis* and *W. chondrophila* were first evidenced by incubating the purified recombinant enzymes overnight at 30°C with 10 mg.mL^-1^ glycogen from rabbit liver (Sigma-Aldrich) and 20 mM inorganic phosphate in 20 mM Tris-HCl buffer (pH 6.8). The reaction products were separated on thin layer chromatography Silica gel 60 W (Merck) using the solvent system butanol/ethanol/water (5/4/3 v/v/v) before spraying orcinol (0.2%)-sulfuric (20%) solution to visualize carbohydrates.

### Maltose-1-phosphate purification

Maltose-1-phosphate was purified from 20 mL of enzymatic reaction with GlgE-EL (1mg GlgE-EL, 10 mg.mL^-1^ potato amylopectin and 20 mM orthophosphate in 20 mM TRIS/HCl pH 6.8, at 30°C, overnight) with several steps. First, size exclusion chromatography on TSK-HW 50 (Toyopearl, 48 x 2.3 cm, flow rate of 1 mL.min^-1^) equilibrated with 1% ammonium acetate was used to remove glycogen. Maltose-1-phosphate was separated from orthophosphate by anion exchange chromatography using Dowex 1 × 8 100-200 mesh (Bio-Rad, 28 x 1.6 cm, acetate form, flow rate of 0.75 mL.min^-1^) in 0.5 M potassium acetate pH 5, then neutralized with ammoniac, and purified from remaining salts using Dowex 50 W X 8 50-100 mesh (Bio-Rad, 10 x 1 cm, H^+^ form) equilibrated with water. Around 10 mg of maltose-1-phosphate were recovered from a reaction mixture of 20 mL, with a yield of approximately 5%. The end product was used for mass spectrometry and NMR analysis. Maltose-1-phosphate was also produced by incubating overnight at 30°C recombinant TreS-Mak protein from *E. lausannensis* with 20 mM ATP, 20 mM maltose, 10 mM MnCl_2_, 125 mM imidazole and 150 mM NaCl. After an anion exchange chromatography as described above, maltose-1-phosphate was purified from remaining maltose and salts using Membra-cell MC30 dialysis membrane against ultrapure water. This purification procedure leads to a better yield (around 8%).

### Proton-NMR analysis of maltose-1-phosphate

Sample was solubilized in D_2_O and placed into a 5mm tube. Spectra were recorded on 9.4T spectrometer (^1^H resonated at 400.33 MHz and ^31^P at 162.10MHz) at 300K with a 5 mm TXI probehead. Used sequences were extracted from Bruker library sequence. Delays and pulses were optimized for this sample.

### MALDI-TOF MS Analysis

P-maltose was analyzed by a MALDI-QIT-TOF Shimadzu AXIMA Resonance mass spectrometer (Shimadzu Europe; Manchester, UK) in the positive mode. The sample was suspended in 20 µL of water. 0.5 µL sample was mixed with 0.5 µL of DHB matrix on a 384-well MALDI plate. DHB matrix solution was prepared by dissolving 10 mg of DHB in 1 mL of a 1:1 solution of water and acetonitrile. The low mode 300 (mass range m/z 250-1300) was used and laser power was set to 100 for 2 shots each in 200 locations per spot.

### Kinetic parameters of GlgE and TreS-Mak of *E. lausannensis*

GlgE activity was monitored quantitatively in the elongation direction by the release of orthophosphate using the Malachite Green Assay Kit (Sigma-Aldrich) following the manufacturers’ instructions. The concentration of released free phosphate was estimated from a standard curve, monitoring the absorbance at 620 nm with Epoch microplate spectrophotometer (Biotek). Kinetic parameters of GlgE-EL were determined in triplicates at 30°C in 15 mM Tris/HCl buffer at pH 6.8.

Saturation plots for maltose-1-phosphate were obtained with 10 mM of maltoheptaose or 10 mg.mL^-1^ of glycogen from bovine liver (Sigma-Aldrich) whereas 2 mM maltose-1-phosphate were used to get saturation plots for maltoheptaose and glycogen from bovine liver. Optimal temperature and pH were assayed with 1 mM maltose-1-phosphate and 5 mM maltoheptaose, respectively in 25 mM Tris/HCl pH 6,8 and at 30°C. Temperature was tested in the range of 15°C to 45°C and pH between 3 and 8.8 with different buffers: 25 mM sodium acetate at pH 3.7, 4.8 and 5.2; 25 mM sodium citrate at pH 3, 4, 5 and 6; 25 mM Tris/HCl at pH 6.8, 7, 7.5, 7.8, 8 and 8,8.

The TreS activity domain of TreS-Mak protein was monitored following the conversion of trehalose into maltose and glucose using Epoch spectrophotometer (Biotek). 15µL of reaction sample were incubated 30 min at 58°C with 30 µL of 100 mM sodium citrate pH 4.6 containing whether 0.4 U of amyloglucosidase from *Aspergillus niger* (Megazyme) or no amyloglucosidase to then quantify the amount of free glucose and maltose after addition of 100 µL of a buffer (500 mM triethanolamine hydrochloride, 3.4 mM NADP^+^, 5 mM MgSO_4_ and 10 mM ATP, pH 7.8). The increase in absorbance at 340 nm after the supplementary addition of 1.2 U of hexokinase and 0.6 U of glucose-6-phosphate dehydrogenase (Megazyme) allowed us to estimate the amount of glucose units from a standard curve. Unless otherwise stated, enzymatic reactions were performed at 30°C, pH 8, with 125 mM imidazole, 150 mM NaCl, 200 mM trehalose and 1 mM MnCl_2_.

The maltokinase activity of TreS-Mak protein was monitored following the amount of nucleoside bi-phosphate released. 20 µL of reaction mixtures were added to 80 µl of pyruvate kinase buffer (75 mM Tris/HCl pH 8.8, 75 mM KCl, 75 mM MgSO4, 2 mM phosphoenolpyruvate, 0.45 mM reduced NADH). The amount of nucleoside diphosphate was estimated from a standard curve of ADP, measuring the decrease in absorbance at 340 nm 30 min after addition of 5 U of L-lactic dehydrogenase (from rabbit muscle, Sigma-Aldrich) and 4 U of pyruvate kinase (from rabbit muscle, Sigma-Aldrich). If not stated otherwise, enzymatic reactions were performed at 30°C, pH 8, with 125 mM imidazole, 150 mM NaCl, 20 mM maltose, 20 mM ATP (or other nucleoside triphosphate) and 10 mM MnCl_2_.

### Chain length distribution analyses

GlgE activity was qualitatively monitored using fluorophore-assisted carbohydrates electrophoresis (FACE). Reactions with 5 mM of malto-oligosaccharides from glucose to maltoheptaose, 1.6 mM M1P and a specific activity of 3.5 nmol orthophosphate produced per minute for GlgE_EL and 1.4 nmol/min for GlgE_WC, were performed in a 100 µL volume at 30°C during 1h and 16h. Reactions were stopped at 95°C for 5 min and supernatants recovered after centrifugation. Samples were dried and resolubilized in 2 µL 1 M sodium cyanoborohydride (Sigma-Aldrich) in THF (tetrahydrofurane) and 2 µL 200 mM ATPS (8-aminopyrene-1,3,6-trisulfonic acid trisodium salt, Sigma-Aldrich) in 15% acetic acid (v/v). Samples were then incubated overnight at 42°C. After addition of 46 µL ultrapure water, samples were again diluted 300 times in ultrapure water prior to injection in a Beckman Coulter PA800-plus Pharmaceutical Analysis System equipped with a laser-induced fluorescence detector. Electrophoresis was performed in a silicon capillary column (inner diameter: 50 µm; outer diameter: 360 µm; length: 60 cm) rinsed and coated with carbohydrate separation gel buffer-N (Beckman Coulter) diluted 3 times in ultrapure water before injection (7 s at 10 kV). Migration was performed at 10 kV during 1 h.

1 mg glycogen from bovine liver (Sigma) and *de novo* polysaccharide produced from overnight incubation of 2 mg maltose-1-phosphate with 30 µg GlgE_EL and 200 µg GlgB_WC were purified by size exclusion chromatography on TSK-HW 50 (Toyopearl, 48 x 2.3 cm, flow rate of 0.5 mL/min) equilibrated with 1% ammonium acetate. Remaining maltose-1-phosphate was dephosphorylated with 10 U of alkaline phosphatase (Sigma-Aldrich) overnight at 30°C and samples were dialyzed using Membra-Cel MC30 dialysis membrane against ultrapure water. The chain length distribution of samples was then analysed following protocol described just above, with slight differences. Prior to APTS labelling, samples were debranched overnight at 42°C in 50 mM sodium acetate pH 4.8 by 2 U of isoamylase from *Pseudomonas* sp. (Megazyme) and 3.5 U of pullulanase M1 from *Klebsiella planticola* (Megazyme), then desalted with AG^®^ 501X8(D) Mixed Bed Resin. Labelled samples were diluted 10 times in ultrapure water before injection.

### Zymogram analysis

7.5% acrylamide-bisacrylamide native gels containing 0.3% glycogen from bovine liver (Sigma-Aldrich) or 0.3% potato starch (w/v) were loaded with 2 µg of crude protein extract or purified recombinant enzyme. Migration was performed in ice-cold TRIS (25 mM) glycine (192 mM) DTT (1 mM) buffer, during 2 h at 120 V and 15 mA per gel, using MiniProtean II (Biorad) electrophoresis system. Gels were then incubated overnight, at room temperature and under agitation, in 10 mL Tris (25 mM) acetate pH 7.5 DTT (0.5 mM), supplemented when stated, with 1 mM maltose-1-phosphate or 20 mM orthophosphate. Gels were rinsed 3 times with ultrapure water prior staining with iodine solution (1% KI, 0.1% I_2_).

### Determination of the apparent molecular weight of GlgE and TreS-Mak

The apparent molecular weight of recombinant GlgE_EL and TreS-Mak_EL were determined using native PAGE and gel filtration. For native PAGE, 5%, 7.5%, 10% and 15% acrylamide: bisacrylamide (37.5 : 1) gels (20 cm x 18.5 cm x 1 mm) were loaded with 6 µg of protein of interest and some standard proteins of known mass: 15 µg carbonic anhydrase (29 kDa), 20 µg ovalbumin (43/86 kDa), 15 µg BSA (66.5/133/266/532 kDa), 15 µg conalbumin (75 kDa), 1.5 µg ferritin (440 kDa) and 25 µg thyroglobuline (669 kDa). Log10 of migration coefficient was plotted against the acrylamide concentration in the gel. Negative slopes were then plotted against molecular weights of standard proteins and the apparent molecular weight of proteins of interest was determined using slope equation. Gel permeation chromatography, Sepharose™ 6 10/300 GL resin (30 cm x 1 cm; GE Healthcare) was equilibrated in PBS buffer (10 mM orthophosphate, 140 mM NaCl, pH 7.4) at 4°C and with a flow rate of 0.3 mL/min. Void volume was determined using Blue Dextran 2000.

Standard proteins used were ribonuclease A (13.7 kDa, 3 mg.mL^-1^), ovalbumin (43 kDa, 4 mg.mL^-1^), aldolase (158 kDa, 4 mg.mL^-1^), ferritin (440 kDa, 0.3 mg.mL^-1^) and thyroglobuline (669 kDa, 5 mg.mL^-1^). All standard proteins used were from GE’s Gel Filtration Low Molecular Weight Kit and GE’S Gel Filtration High Molecular Weight Kit (GE Healthcare), except for the BSA (Sigma-Aldrich).

## Results

### Two different glycogen metabolic pathways identified in the *Chlamydiae* phylum

To gain insight into Chlamydiae’s glycogen metabolism, we analyzed 220 genomes from 47 different chlamydial species. As illustrated in **figure 1A**, the synthesis of linear chains of α-1,4 glucose involves both ADP-glucose pyrophosphorylase (GlgC) and glycogen synthase (GlgA) activities in the GlgC-pathway while GlgE-pathway relies on trehalose synthase (TreS), maltokinase (Mak) and maltosyl-1 phosphate transferase (GlgE). The formation of α-1,6 linkages (i.e. branching points) and glycogen degradation are catalyzed by a set of similar enzymes in both pathways that include glycogen branching enzyme isoforms (GlgB/GlgB2) and glycogen phosphorylases isoforms (GlgP /GlgP2), glycogen debranching enzymes (GlgX) and α-1,4 glucanotransferase (MalQ). The genomic database used in this study (https://chlamdb.ch) includes genomes from both cultured and uncultured Chlamydiae species that cover the diversity of the chlamydiae phylum (**figure 1B**). Comparative genomics clearly underlined the high prevalence of a complete GlgC-path in most *Chlamydiales,* including all members of the *Chlamydiaceae* family, which undergoes massive genome reduction (identified by the letter “d” on **figure 1B**) as well as in in the most deeply branching families such as candidatus *Pelagichlamycidiaceae* (“a”) and candidatus *Parilichlamydiaceae* (“b”). We noticed that the *glg* genes are at least 10 kbp apart with a notable exception for *glgP* and *glgC*, which are mostly separated by one or two genes. It should be stressed out that the gaps in glycogen metabolism pathways of several uncultivated chlamydiae likely reflect the fact that many of those genomes are incomplete genomes derived from metagenomic studies (see percentages in brackets in **figure 1B**). Considering that the GlgC-pathway is highly conserved in nearly all sequenced genomes of the phylum, missing genes probably reflect missing data rather than gene losses. It is interesting to note that there is uncertainty about the presence of *glgC* gene in Candidatus *Enkichlamydia* genome (“j”), a complete set of glycogen metabolizing enzymes were recovered expect for gene encoding for ADP-glucose pyrophospharylase (glgC). This gene is missing from 6 independent draft genomes estimated to be 71% to 97% complete, suggesting either the loss of *glgC* gene or that *glgC* gene is located in a particular genomic region (e.g. next to repeated sequences) that systematically led to its absence from genome assemblies. Another unexpected result concerns both *Waddliaceae* (“l”) and *Criblamydiaceae* (“m”) families that encompass *Waddlia chondrophila*, *Estrella lausannensis* and *Criblamydia sequanensis* species. Genomic rearrangements caused a sequence of events leading to (i) the deletion of both *glgC* and *glgP* genes (ii) the fusion of *glgA* with *glgB* gene (iii), the insertion of *glgP2* gene encoding glycogen phosphorylase isoform at the vicinity of *malQ* gene. It should be stressed out that a homolog of *glgP2* gene has also been identified on the plasmids of *S. nevegensis* and *P. naegleriophila*. In *W. chondrophila*, another insertion of *glgP2* occurred downstream to the GlgE operon, which may be correlated with partial deletion of *glgP2* at the vicinity of *malQ* (**Figure 1B**). The parsimonious interpretation of *glgC* and *glgP* deletion and *glgAglgB* fusion is that a single deletion event led to the loss of DNA fragment bearing *glgP* and *glgC* genes between *glgA* and *glgB*. Despite many variations, we did not observe such configuration in the chlamydial genome analyzed (**S2 Table**). More remarkably, genomic rearrangements are associated with a novel glycogen pathway based on GlgE operon described in mycobacteria and also observed in *Prototochlamydia naegleriophila* and *Protochlamydia phocaeensis* (syn. *Parachlamydia* C2). All three genes are clustered in the classical unfused *glgE-treSmak-glgB2* operon arrangement in *Waddliaceae* and *Criblamydiaceae*, while the *glgB2* gene is missing in the *Parachlamydiaceae* operons (**Figure 1B**). The occurrence of GlgE pathway restricted to *Parachlamydiaceae*, *Waddliaceae* and *Criblamydiaceae* families arises questions about its origin in Chlamydiales. To get some insight on this issue, phylogenetic trees of TreS-Mak and GlgE have been inferred using the phylobayes method (**Figure 2**). The GlgE phylogeny shows that even if the *Chlamydiae* sequences are split into two with *W. chondrophila* on one side and the other sequences on the other side, which reflects likely lateral gene transfer events with other bacteria, chlamydial *glgE* sequences might still be monophyletic since the only strongly supported node (marked as red star) with a posterior probability (pp) higher than 0.95 (pp = 0.99) unifies all chlamydiae sequences, which has also been confirmed using LG4X model (Data not shown) (**Figure 2A**). The phylogeny analysis highlights that GlgE sequences can be classified into classes I and II, comprising *Chlamydiales* and *Actinomycetales* (i.e. mycobacteria, Streptomycetes), respectively. For Tres-Mak phylogeny, chlamydial Tres-Mak sequences cluster together, suggesting a common origin, however with a low statistical support (pp=0.93). Although the origin of GlgE operon cannot be pinpointed in our phylogenetic analysis, conceivable scenarios are that either i) GlgE operon reflects vestigial metabolic function of the ancestral chlamydiae and then has been lost in most families or ii) this operon was acquired by lateral gene transfer event from a member of PVC phylum by the common ancestor of *Parachlamydiaceae*, *Waddliaceae* and *Criblamydiaceae* families.

**Figure 1:**
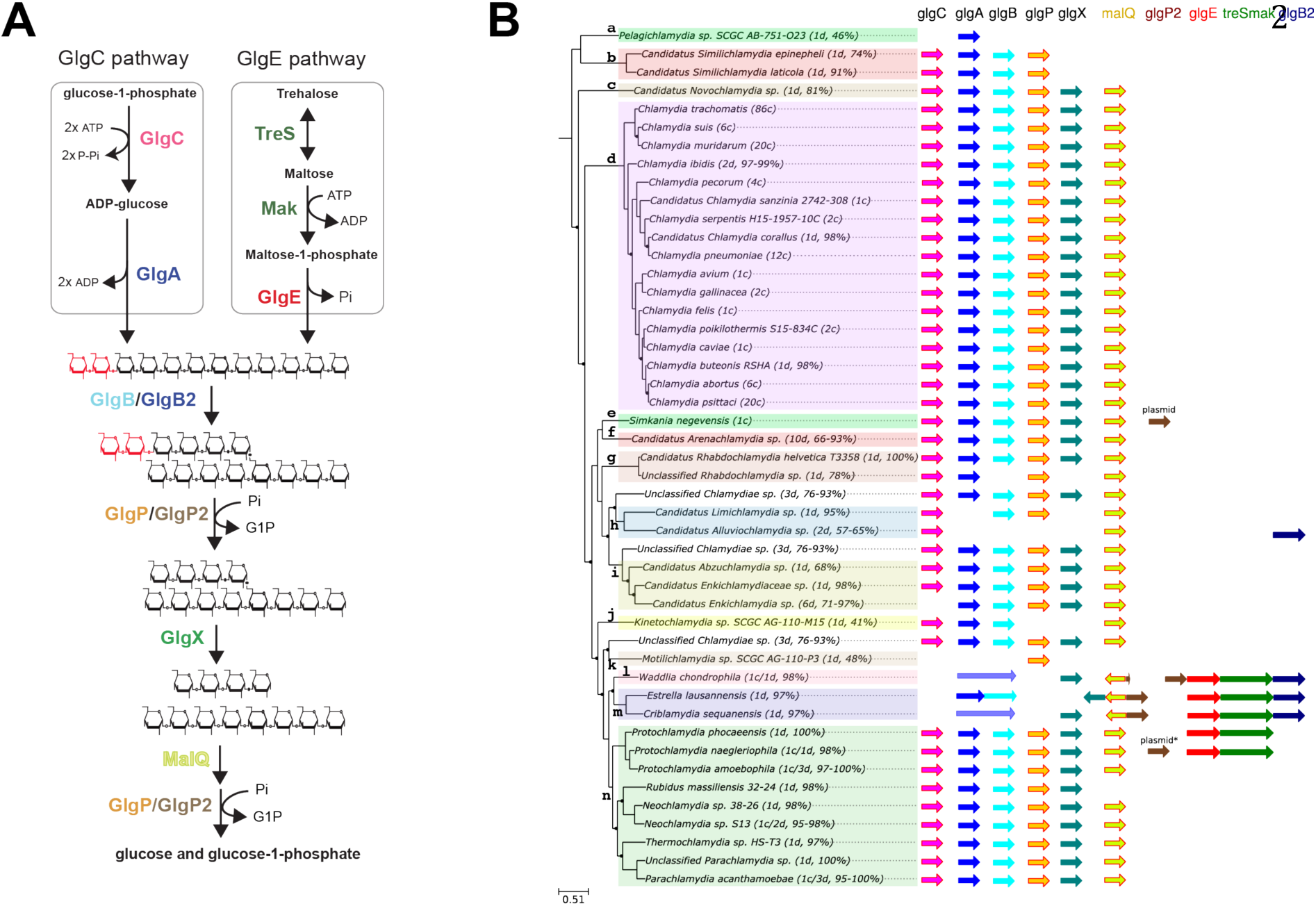
Comparative genomic analysis of glycogen metabolizing genes among *chlamydiae* phylum. **A**. GlgC-and GlgE-paths represent the main routes of glycogen biosynthesis in prokaryotes. The formation of linear chains of glucosyl units joined by α-1,4 linkages depends on the coupled actions of ADP-glucose pyrophosphorylase (GlgC)/glycogen synthase (GlgA) activities in GlgC-path whereas it relies on the combined actions of trehalose synthase (TreS)/maltokinase (Mak)/ maltosyltransferase (GlgE) in GlgE-path. The iteration of glucan synthesis and branching reactions catalyzed by branching enzyme isoforms (GlgB and glgB2) generate a branched polysaccharide. Both α-1,4 and α-1,6 glucosidic linkages are catabolized through synergic actions of glycogen phosphorylase isoforms (GlgP and GlgP2), debranching enzyme (GlgX) and a-1,4 glucanotransferase (MalQ) into glucose-1-phosphate and glucose **B** Phylogenic tree of cultured and unculturedchlamydiae.. For each species of the families: a, Ca. *Pelagichlamydiaceae*; b, Ca. *Paralichlamydiaceae*; c, Ca. *Novochlamydiacae*; d, *Chlamydiaceae*; e, *Simkaniaceae*; f, Ca. *Arenachlamydiaceae*; g, *Rhabdochlamydiaceae*; h, Ca. *Limichlamydiaceae*; i, Ca. *Enkichlamydiaceae*; j, Ca. *Kinetochlamydiaceae*; k, Ca. *Motilichlamydiaceae*; l, *Waddliaceae*; m, *Criblamydiaceae*; n, *Parachlamydiaceae*, the number of draft (d) or complete (c) genomes and genome completeness expressed in percentage are indicated between brackets. Homologous genes of the GlgC- and GlgE-pathways are symbolized with colored arrows. The glgP2 gene was identified on the plasmid of *S. negevensis* and is also present in one of the two *P. neagleriophila* genome available.

**Figure 2:**
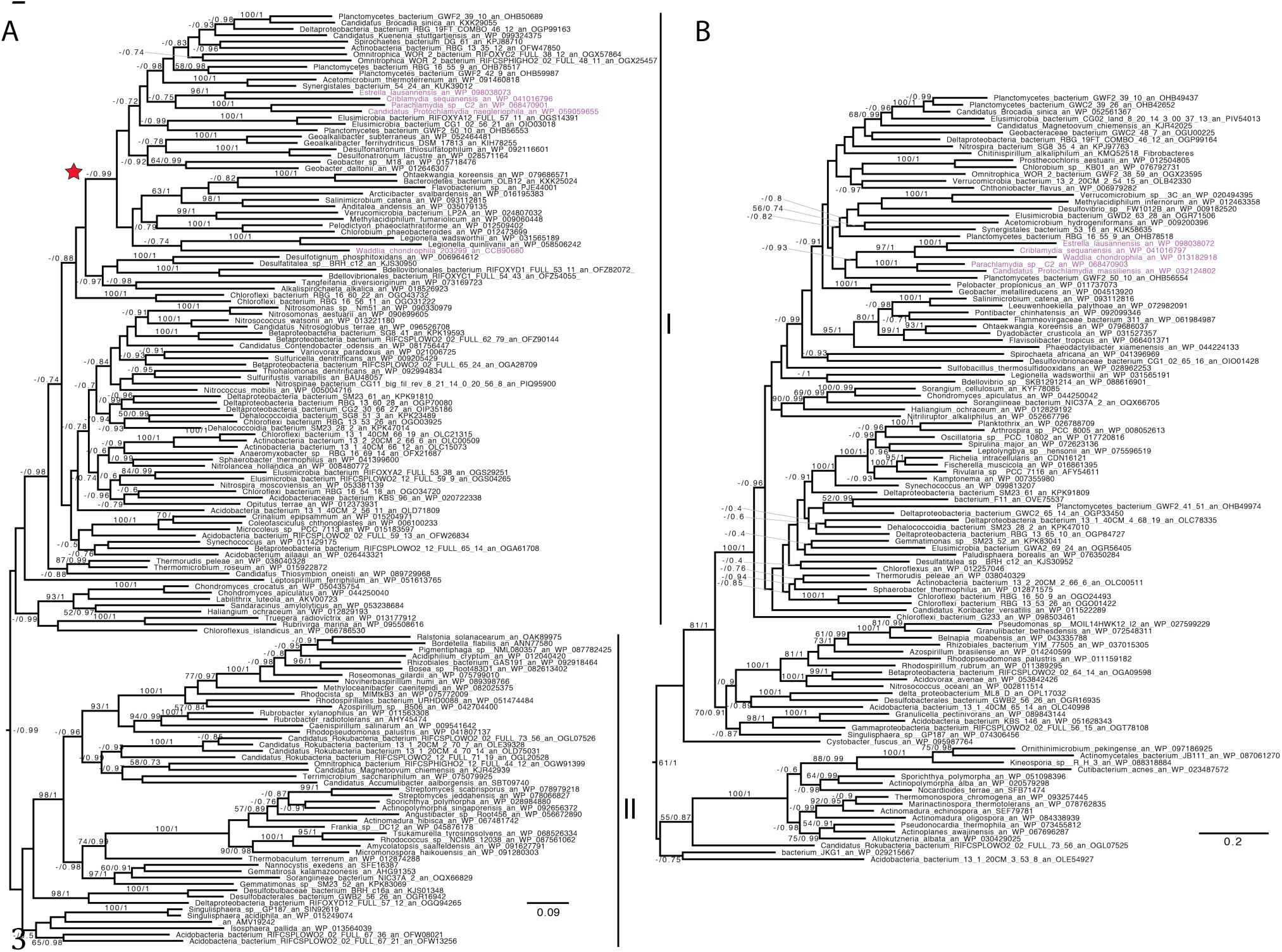
Phylogenetic analysis of GlgE (A) and TreS-Mak (B). Trees displayed were performed with Phylobayes under the C20+poisson model. We then mapped onto the nodes ML boostrap values obtained from 100 bootstrap repetitions with LG4X model (left) and Bayesian posterior probabilities (right). Bootstrap values >50% are shown, while only posterior probabilities >0.6 are shown. The trees are midpoint rooted. The scale bar shows the inferred number of amino acid substitutions per site. The *Chlamydiales* are in purple.

### Classical GlgC-pathway is not functional in *E. lausannensis* and *W. chondrophila*

To further investigate whether his-tagged recombinant proteins GlgA-GlgB of *E. lausannensis* and *W. chondrophila* are functional, glycogen synthase activities at the N-terminus domain were assayed by measuring the incorporation of labeled ^14^C-glucosyl moiety from ADP- or UDP-^14^C-glucose onto glycogen and by performing a specific non-denaturing PAGE or zymogram to visualize glycogen synthase activities. After separation on native-PAGE containing glycogen, recombinant proteins were incubated in the presence of 1.2 mM ADP-glucose or UDP-glucose, glycogen synthase activities are visualized as dark activity bands after soaking gels in iodine solution (**Figure 3**). Enzymatic assays and zymogram analyses show that the glycogen synthase domain of the chimeric GlgA-GlgB of *W. chondrophila* (hereafter GlgA-GlgB-WC) is functional but highly specific for ADP-glucose (0.70 nmol of incorporated glucose. min^-1^.mg^-1^) and has little to no activity using UDP-glucose as substrate. As predicted, the activity of the truncated glycogen synthase in *E. lausannensis* was not detected on activity gels or during enzymatic assays (**S1A Figure**).

**Figure 3:**
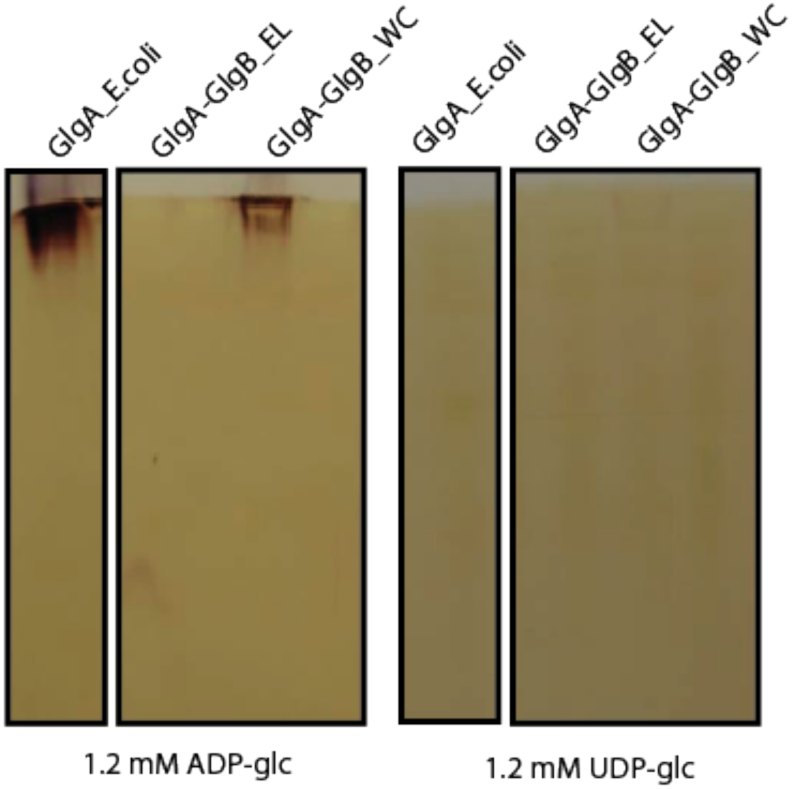
Zymogram analysis of glycogen synthase activities. **A**. Total crude extracts of the recombinant proteins of GlgA of *Escherichia coli* (GlgA_E.coli), GlgA-GlgB of *E. lausannensis* (GlgA-GlgB_EL) and *W. chondrophila* (GlgA-GlgB_WC) were separated by native PAGE containing 0.6% (w/v) glycogen. The native gels were then incubated with 1.2 mM ADP-glc or 1.2 mM UDP-glc. Glycogen synthase activities are seen after iodine staining as dark bands.

We further investigated whether the branching activity domain at the carboxyl terminal of chimeric protein GlgA-GlgB of *W. chondrophila* (GlgA-GlgB-WC) was functional. To check this, the same chimeric GlgA-GlgB-WC sample previously analyzed was incubated with ADP-glucose (3 mM) and maltoheptaose (10 mg.mL^-1^) overnight. Subsequently, the appearance of branching point (*i.e* α-1,6 linkages) onto growing linear glucans can be specifically observed by the resonance of protons onto carbon 6 at 4.9 ppm using proton-NMR analysis. However as depicted on **S1C figure**, we did not observe any signal, suggesting that branching enzyme activity domain is not functional despite an active glycogen synthase domain. This result is consistent with several reports indicating that the amino-acid length at the N-terminus of branching enzyme affects its catalytic properties [36, 37]. In regards to this information, the glycogen synthase domain extension located at the N-terminus prevents probably the branching enzyme activity of GlgA-GlgB. Thus α-1,6 linkages or branching points are likely to be the result of GlgB2 isoform activity found in both instances. Altogether, these data strongly suggest that the classical GlgC-pathway is not functional in both *Waddliaceae* and *Criblamydiaceae* families.

### GlgE-like genes of *E. lausannensis* and *W. chondrophila* encode α-maltose 1-phosphate: 1,4-α-D-glucan 4-α-D-maltosyl transferase

Based on phylogenetic analysis of GlgE, both GlgE of mycobacteria (*Actinobacteria*) and *Chlamydiales* are phylogenetically distant (**Figure 2A**). GlgE of *M. tuberculosis* displays 43% to 40% of identity with GlgE-like sequences of *E. lausannensis* and *W. chondrophila*, respectively. Because GlgE activity belongs to the large and diversified Glycosyl Hydrolase 13 family consisting of carbohydrate active enzymes with quite diverse activities such as α-amylases, branching enzymes, debranching enzymes [38], we undertook to demonstrate that these enzymes displayed catalytic properties similar to those previously described for GlgE of mycobacteria. Histidine-tagged recombinant proteins of GlgE of *Estrella lausannensis* (hereafter GlgE-EL) and *Waddlia chondrophila* (hereafter GlgE-WC) were expressed and further characterized (**S2 Figure**). As described in previous studies, GlgE of *Mycobacteria* mediates the reversible reaction consisting into the release of maltose-1-phosphate in the presence of orthophosphate and α-glucan polysaccharide. Both GlgE-EL and GlgE-WC were incubated in presence of glycogen from rabbit liver and orthophosphate. After overnight incubation, reaction products were analyzed on thin layer chromatography and sprayed with oricinol-sulfuric acid (**Figure 4A**). A fast migration product capable of interacting with oricinol sulfuric acid was clearly synthesized in crude extract (CE), in washing # 3 (W3) and in the purified enzyme fraction (E1) of GlgE-EL sample. A barely visible product is only observed in the purified fraction (E1) of GlgE-WC. To further characterize this material, time course analysis of phosphatase alkaline (PAL) treatment was performed on the reaction product suspected to be M1P obtained from sample E1 of GlgE-EL. After 180 min of incubation, the initial product is completely converted into a compound with a similar mobility than maltose (DP2) (**Figure 4B**). The compound produced by GlgE-EL in presence of glycogen and orthophosphate were further purified through different chromatography steps and subjected to mass spectrometry and proton-NMR analyses (**Figures 4C** and **4D**). The combination of these approaches confirms that GlgE of *E. lausannensis* as well as *W. chondrophila* (**S3 Figure**) catalyzes the formation of a compound of 422 Da corresponding to α-maltose-1-phosphate. In order to carry out enzymatic characterization of GlgE activities, identical purification processes were scaled up to purify enough M1P, free of inorganic phosphate and glucan.

**Figure 4:**
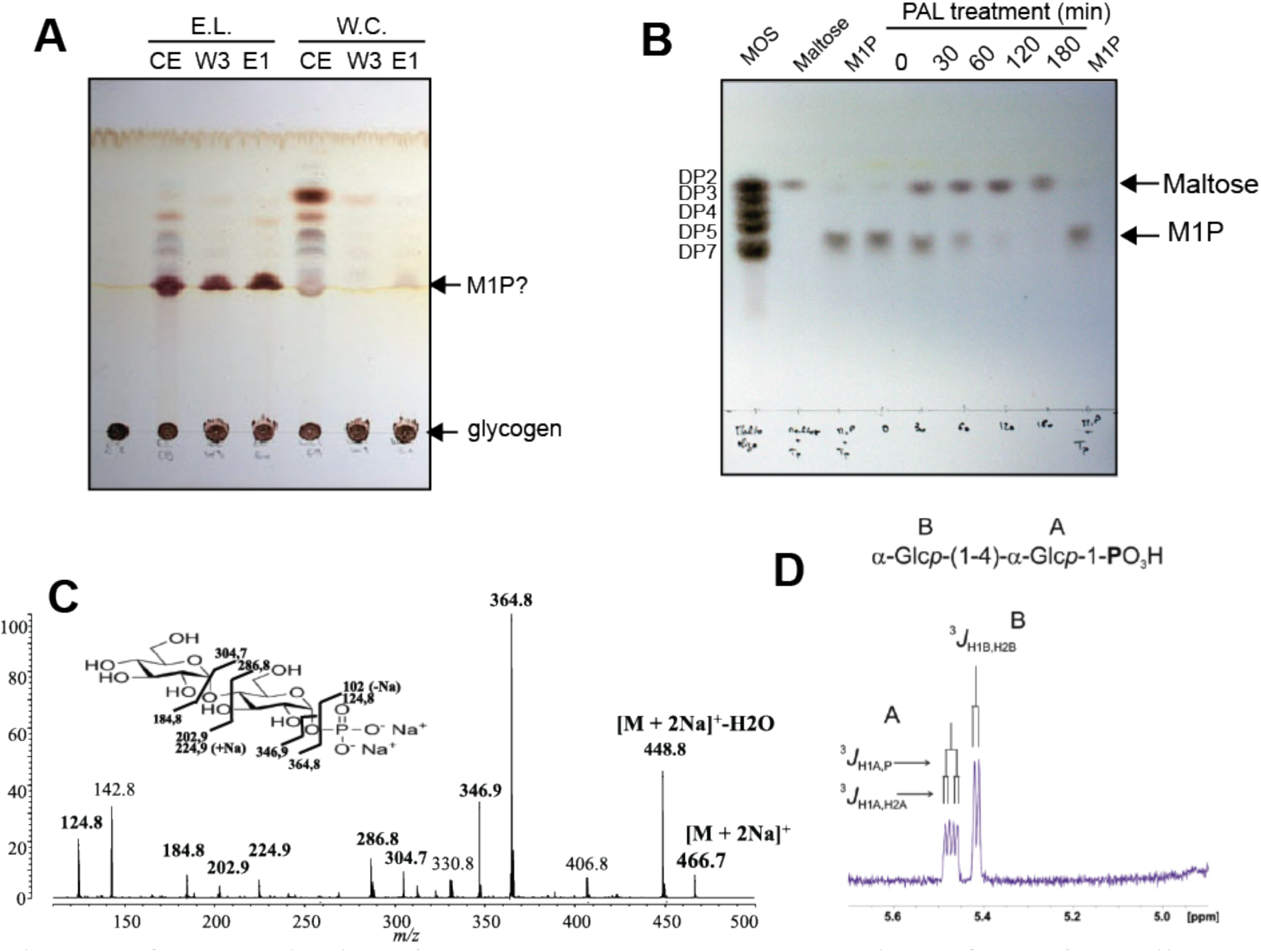
Characterization of compounds released by recombinant GlgE of *Estrella lausannensis*. **A**, both histidine-tagged recombinant GlgE-EL and GlgE-WC proteins were purified and incubated in presence of glycogen and inorganic phosphate. The overnight reaction products from crude extract (CE), third washing step (W3) purified enzymes (E1) were subjected to thin layer chromatography analysis. Orcinol-sulfuric spray reveals a significant production of M1P with recombinant GlgE-EL, which is less visible with recombinant GlgE-WC. **B**, after one purification step consisting in the removal of glycogen by size exclusion chromatography, M1P was incubated in presence of phosphatase alkaline (PAL). Times course analysis (0 to 180 minutes) shows a complete conversion of M1P into maltose. **C**, MS-MS sequencing profile of M1P. The molecular ion [M + 2Na]^+^ at m/z 466,7 corresponding to M1P + 2 sodium was fractionated in different ions. Peak assignments were determined according to panel incrusted in C. **D**, part of 1D-^1^H-NMR spectrum of maltoside-1-phosphate. α-anomer configuration of both glucosyl residues were characterized by their typical homonuclear vicinal coupling constants (^3^*J*_H1A,H2A_ and ^3^*J*_H1B,H2B_) with values of 3.5 Hz and 3.8 Hz respectively. A supplementary coupling constant was observed for α-anomeric proton of residue A as shown the presence of the characteristic doublet at 5.47 ppm. This supplementary coupling constant is due to the heteronuclear vicinal correlation (^3^*J*_H1A,P_) between anomeric proton of residue A and phosphorus atom of a phosphate group, indicating that phosphate group was undoubtedly *O*-linked on the first carbon of the terminal reducing glucosyl unit A. The value of this ^3^*J*_H1A,P_ was measured to 7.1Hz.

### Kinetic parameters of GlgE activity of *E. lausannensis* in the biosynthetic direction

Because the his-tagged recombinant GlgE-WC expresses very poorly and the specific activity of GlgE-WC was ten times lower than GlgE-EL, kinetic parameters were determined in the synthesis direction *i.e.* the transfer (amount) of maltosyl moieties onto non-reducing ends of glucan chains, exclusively for GlgE-EL. Transfer reactions are associated with the release of inorganic phosphate that can be easily monitored through sensitive malachite green assay. Thus, under variable M1P concentrations and using fixed concentrations of glycogen or maltoheptaose, the GlgE-EL activity displays allosteric behavior indicating a positive cooperativity, which has been corroborated with Hill coefficients that were above 1 (**Figures 5 A** and **5 B**). In agreement with this, the molecular weight of native GlgE-EL determined either by size exclusion chromatography or by native-PAGE containing different acrylamide concentrations (5%; 7.5 %; 10 % and 12.5 %) indicates an apparent molecular weight of 140 to 180 kD respectively corresponding to the formation of dimer species while no monomer species of 75 kD were observed (**Figure 5E**). The enzyme exhibited S_0.5_ values for M1P that vary from 0.16±0.01 mM to 0.33±0.02 mM if DP7 and glycogen are glucan acceptors, respectively. However, using M1P at saturating concentration, GlgE-EL displays Michaelis kinetics (n_H_ close to 1) indicating a non-cooperative reaction (**figures 5C** and **5 D**). In such experimental conditions, the apparent K_m_ values for glycogen and DP7, 2.5 ± 0.2 mg.mL^-1^ and 3.1 ± 0.2 mM, respectively were similar to the apparent Km value of glycogen synthase (GlgA) that synthesizes α-1,4 linkages from ADP-glucose [39].

**Figure 5:**
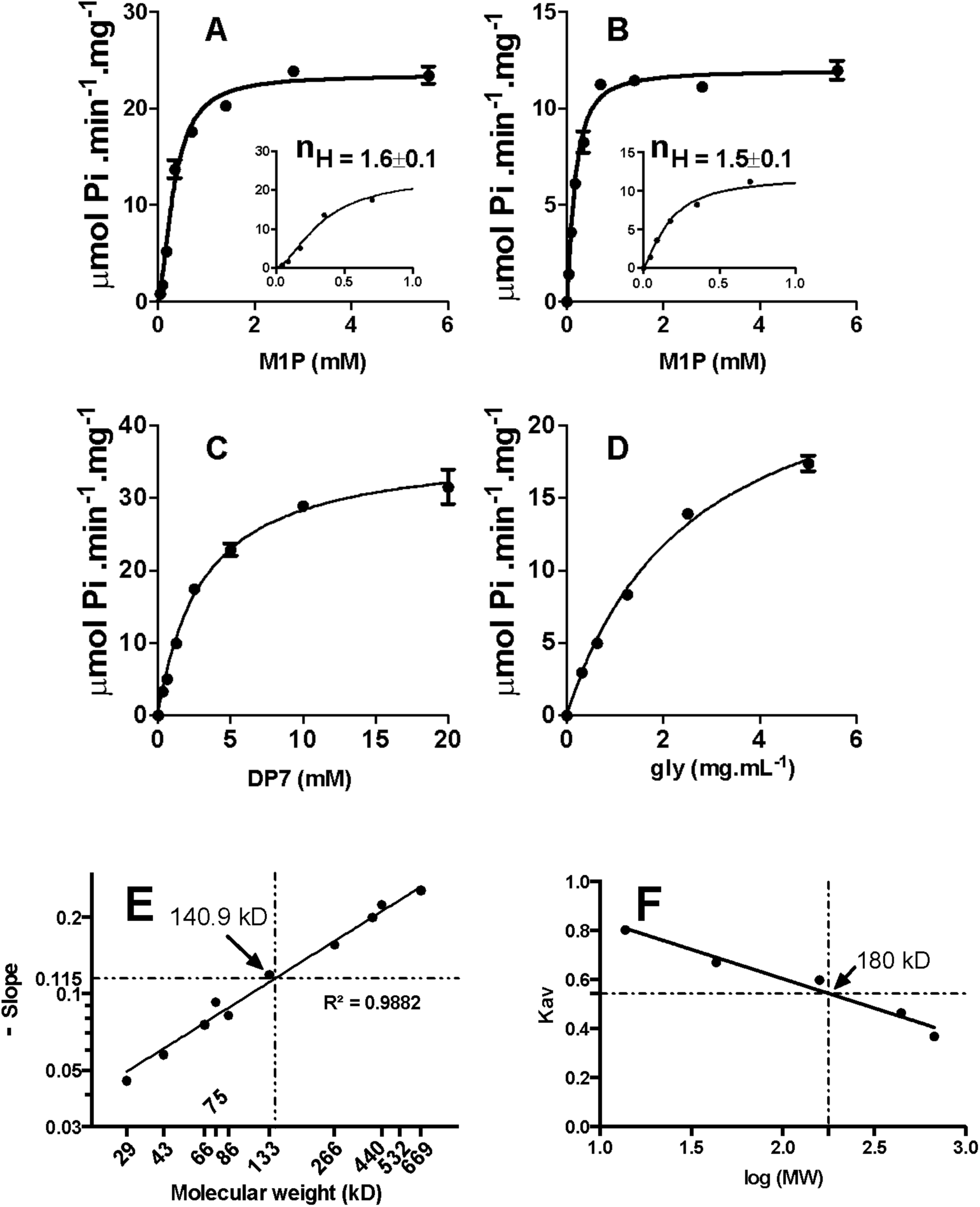
Kinetic parameter of recombinant GlgE-EL. GlgE activity was assayed spectrophotometrically by monitoring the release of inorganic orthophosphate (Pi). Kinetic constants were determined in triplicate. M1P saturation plots for GlgE-EL were determined in the presence of 10 mM of maltoheptaose (DP7) (**A**) or 10 mg.mL^-1^ of glycogen (**B**). At low M1P concentrations (panels), GlgE-EL activity behaves as allosteric enzyme with Hill coefficients (n_H_) of 1.6 and 1.5, respectively (fit shown as the solid line giving r^2^=0.98). The S_0.5_ (M1P) values for GlgE-EL were determined at 0.33±0.02 mM and 0.16±0.01 mM in the presence of DP7 and glycogen, respectively. In the presence of 2 mM M1P, both DP7 (**C**) and glycogen (**D**) saturation plots are conform to the Michaelis-Menten behavior (n_H_ close to 1) with Km values of 3.1±0.2 mM and 2.5±0.2 mg.mL^-1^, respectively. The apparent molecular weight of GlgE-EL was determined by native PAGE (**E**) and size exclusion chromatography (Superose 6 Increase GL 10/300) (**F**) at 140.9 kDa and 180 kDa respectively suggesting a dimer of GlgE (76 kD).

### *De novo* glycogen synthesis: GlgE activity enables the initiation and elongation of glucan

Contrarily to eukaryotic glycogen synthase, prokaryotic glycogen synthase (GlgA) does not require the presence of a short α-1,4 glucan or primer to initiate glycogen biosynthesis [9]. In absence of GlgA and GlgC activity in *E. lausannensis* and in the absence of GlgC and thus of ADP-glucose supply in *W. chondrophila*, this raised the question of the ability of GlgE activities to substitute for GlgA with respect to the priming of glycogen biosynthesis. To establish whether GlgE activities are able to prime glucan synthesis, both his-tagged GlgE-EL (3.51 nmol of Pi released.min^-1^) and GlgE-WC (1.38 nmol of Pi released.min^-1^) were incubated with 1.6 mM M1P and in the presence of 5mM of various glucan chains with a degree of polymerization (DP) of 1 to 7. Identical incubation experiments were conducted with GlgE recombinant proteins except M1P was omitted in order to appreciate α-1,4 glucanotransferase or disproportionnating activity (**Figure 6** and **S4, S5 Figures**). After incubation, the reduced-ends of glucan chains were labeled with fluorescent charged probe (APTS) and separated according to their degree of polymerization by capillary electrophoresis. We noticed that the C1 phosphate group prevented the labeling of M1P with fluorescent probe.

**Figure 6:**
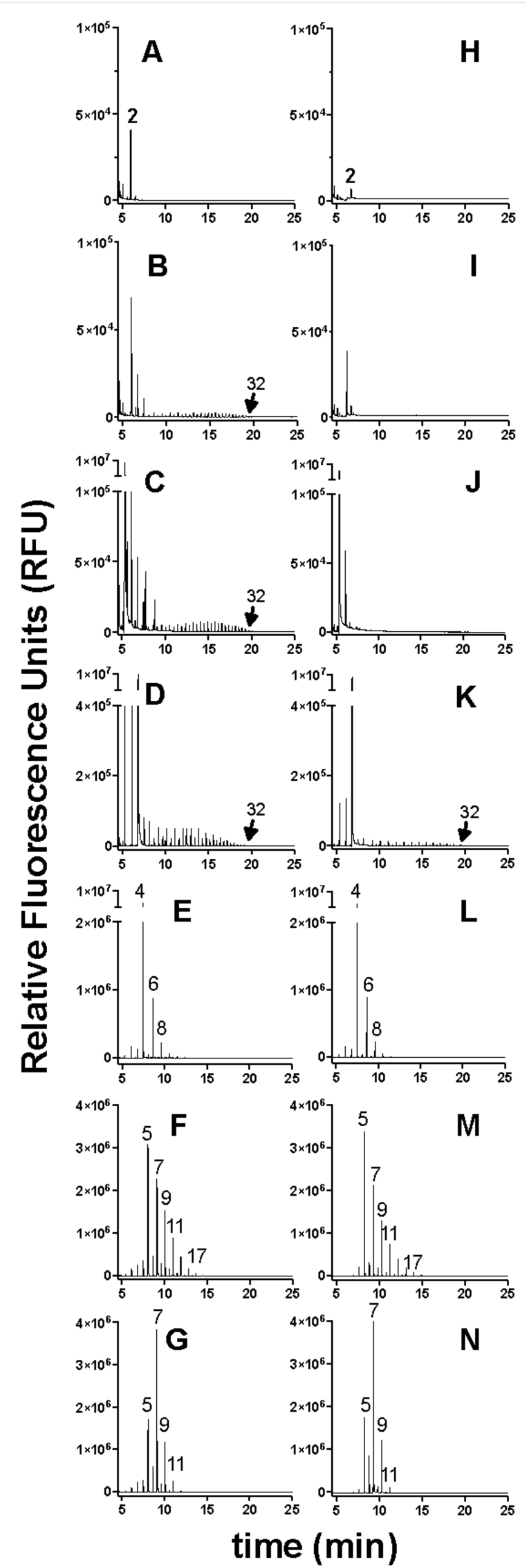
FACE analyses of enzymatic reaction products of GlgE activity of *W. chondrophila*. (A to G) and *E. lausannensis* (H to N). Spontaneous dephosphorylation of M1P during overnight incubation was estimated by incubating denatured GlgE enzymes in buffer containing 1.6 mM M1P (A and H). The transfer of maltosyl moieties from M1P at 1.6 mM onto non-reducing ends of glucan acceptors (5 mM) were determined in absence of glucan acceptor (B, I) or in presence of glucose, (C, J)), maltotriose (D, K), maltotetraose (E, L) and maltoheptaose (F,M). α-1,4-glucanotransferase activities of GlgE were determined by incubating 5 mM of maltoheptaose without maltose-1-phosphate (G, N). Numbers on the top of fluorescence peaks represent the degree of polymerizations of glucan chains.

Nevertheless, the level of maltose released from M1P due to the spontaneous dephosphorylation during the experiment was appreciated by performing incubations with denatured enzymes (**Figures 6A and 6H**). Incubation experiments show that both GlgE activities possess either an α-1,4 glucanotransferase or maltosyltransferase activities depending on the presence of M1P. When M1P is omitted, GlgE activities harbor an α-1,4 glucanotransferase activity exclusively with glucans composed of six or seven glucose units (DP6 or DP7). Interestingly, after one hour or overnight incubation, DP6 or DP7 are disproportionated with one or two maltosyl moieties leading to the release of shorter (DPn-2) and longer glucans (DPn+2) (**Figures 6G and 6N** and **S4, S5 Figures**). The limited number of transfer reactions emphasizes probably a side reaction of GlgE activities. The α-glucanotransferase activity can be also appreciated on native-PAGE containing glycogen. Chain length modification of external glucan chains of glycogen results in increase of iodine interactions visualized as brownish activity band (**S6A Figure**). After one hour of incubation (**S4, S5 Figures**), both GlgE activities enable the transfer the maltosyl moiety of M1P onto the glucan primer with a DP ≥ 3 (**S4, S5 Figures**). Interestingly, for a longer period of incubation time, both GlgE activities switch to either a processive or a distributive activity mode depending on the initial degree of polymerization of glucan primer. For instance, in the presence of maltose (DP2) or maltotriose (DP3) both GlgE-EL and GlgE-WC undergo processive elongation activities, which consist of the synthesis of very long glucan chains, up to 32 glucose residues. In contrast, when both GlgE activities are incubated in presence of glucan primers with DP ≥4, the latter add and immediately release a glucan primer (DP) with an increment of two glucose moieties (DPn+2) that leads to distributive elongation behavior. The mechanism underlying the switch between processive and distributive elongation activities reflects probably a competition of glucan primers for the glucan binding site in the vicinity of the catalytic domain. Thus, we can hypothesize that the low affinity of short glucan primers (DP<4) for glucan binding site favors probably iterative transferase reactions onto the same acceptor glucan (i.e. processive mode) resulting in the synthesis of long glucan chains whereas glucan primers with DP≥ 4 compete strongly for the binding site leading to a distributive mode. The discrepancy between GlgE-EL and GlgE-WC to synthesize long glucan chains in the absence (**Figures 6B, 6I**) or in the presence of glucose (DP1) (**Figures 6C, 6J**) might be explained by a higher amount of free maltose observed in denatured GlgE-WC samples (**Figure 6A**) by comparison to denatured GlgE-EL samples (**Figure 6H**). Despite having taken all precautious (same M1P preparation, buffer pH7), spontaneous dephosphorylation of M1P occurred more significantly in GlgE-WC samples. We therefore conclude that initial traces of maltose in GlgE-WC samples facilitate the synthesis of long glucan chains in the absence (**Figure 6B**) or in the presence of glucose (DP1) (**Figure 6C**). To test this hypothesis, crude extract (CE) and purified GlgE proteins (E1) of *E. lausannensis* were loaded onto non-denaturing polyacrylamide electrophoresis (native-PAGE). After migration, slices of polyacrylamide gel were incubated overnight in buffers containing 0 mM (control) or 2 mM M1P (**Figure 7A**). The synthesis of long glucan chains with DP> 15 (minimum number of glucose units required for detection through interaction with iodine molecules) are detected by soaking the gel in iodine solution. As depicted in **Figure 7A**, the synthesis of glucan chains catalyzed by GlgE-EL appears exclusively as dark-blue activity bands inside native-PAGE incubated with 2 mM M1P and not in the absence of M1P. Altogether, these results suggest that GlgE activities are able to synthesize *de novo* a sufficient amount of long linear glucans from maltose-1-phosphate. We cannot exclude the role of maltose in the initiation process of glucan synthesis as glucan acceptor since spontaneous dephosphorylation of M1P is unavoidable. We further carried out a series of experiments that consisted to synthesize *in vitro* high molecular branched glucans by incubating both recombinant glycogen branching enzyme of *W. chondrophila* (GlgB-WC: **S6B Figure**) and GlgE-EL in the presence of M1P. After overnight incubation, the appearance of α-1,6 linkages or branching points were directly measured by subjecting incubation product on proton-NMR analysis (**Figure 7B**). In comparison with M1P and glycogen as controls, proton-NMR spectrum of incubation products show a typical profile of glycogen-like with signals at 5.6 ppm and 4.9 ppm of proton involved in a-1,4 and a-1,6 linkages.

**Figure 7:**
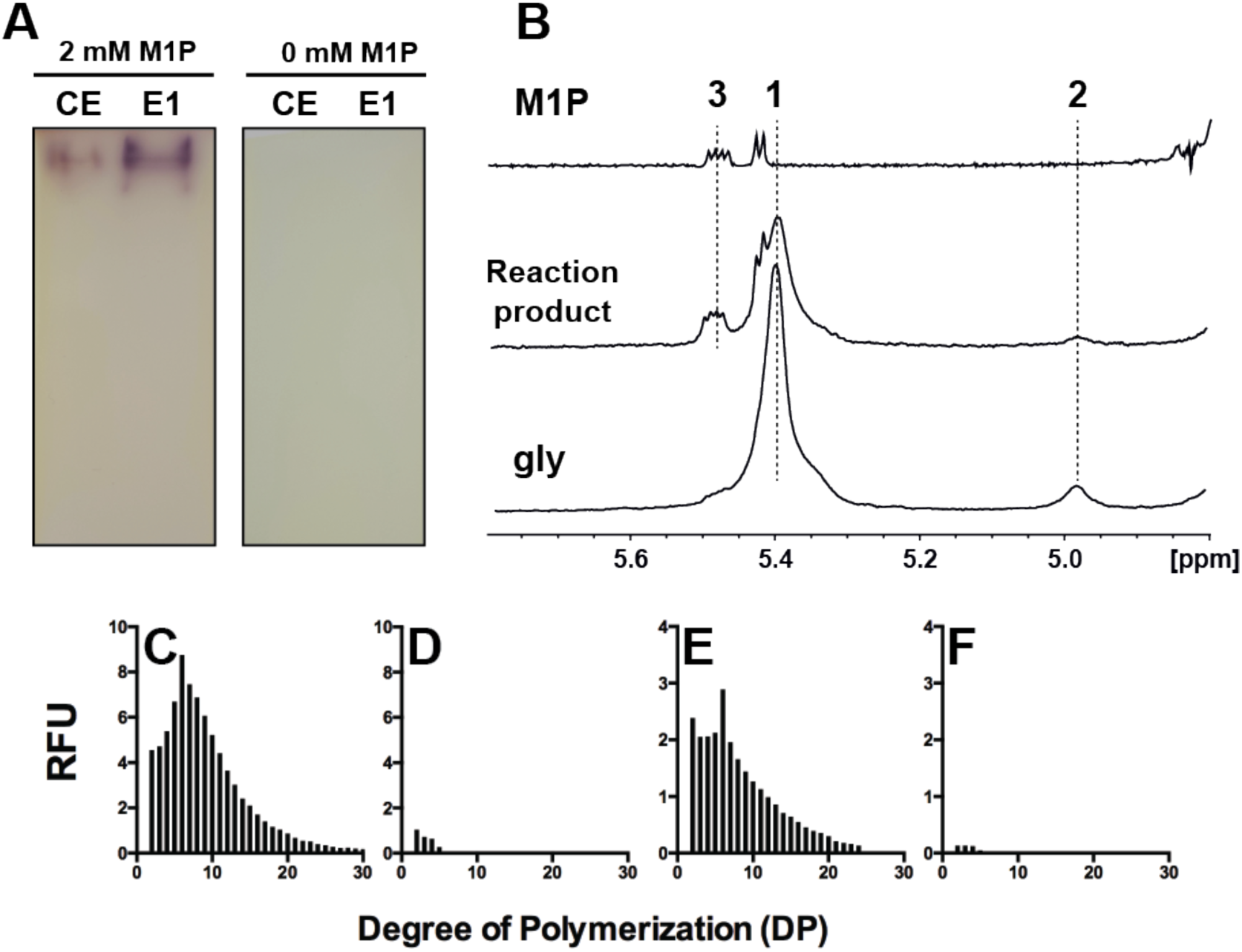
*De novo* synthesis of branched polysaccharides. **A**. Recombinant GlgE activity of *Estrella lausannensis* from crude extract of *E. coli* (CE) and purified on nickel affinity column (E1) were loaded on non-denaturing polyacrylamide gel. After migration, slices of native-PAGE were incubated in TRIS/acetate buffer containing 2 mM of maltose-1-phosphate (M1P) over 16 hours at 25°C. The synthesis of *de novo* glucan chains is visualized as dark-blue bands due to the formation of glucan-iodine complexes. Similar *in vitro* experiments were conducted by adding GlgB activity of *W. chondrophila* to a TRIS/acetate buffer containing GlgE activity and 2 mM of M1P. After overnight incubation, reaction mixture was subjected to ^1^H-NMR analysis. **B.** Part of ^1^H-NMR spectra of maltose-1-phosphate (M1P), non-purified reaction mixture and glycogen (gly) from bovine liver in D_2_O. Peak #1 (5.45 to 5.3 ppm) and peak #2 (4.98 ppm) represent the signals of protons involved respectively in α-1,4 and α-1,6 linkages while peak #3 (5.47 ppm) represents the characteristic doublet of doublet signals of α-anomeric proton located on C1 of maltose-1-phosphate. The appearances of peak #2 and peak #3 in incubation product indicate the formation of branched polysaccharide composed of α-1,4 and α-1,6 linkages. The presence of peak #3 suggests that M1P was not completely polymerized by GlgE activity of EL. α-Polysaccharides were then purified (see materials and methods) and incubated with a commercial isoamylase type debranching enzyme. After overnight incubation, the linear glucan chains released from α-polysaccharides (**C**) and glycogen from bovine liver used as reference (**D**) were separated according to the degree of polymerization by capillary electrophoresis coupled with a fluorescent labeling of reduced-ends. As control, α-polysaccharide (**D**) and glycogen (**F**) samples were directly labeled and analyzed by capillary electrophoresis in order to estimate the content of free-linear glucan chains.

This branched polysaccharide material was further purified and incubated with a commercial isoamylase type debranching enzyme (Megazyme) that cleaves off α-1,6 linkages or branching points. Released linear glucan chains were labeled with APTS and separated according to the degree of polymerization by capillary electrophoresis. The chain length distribution (CLD) of synthesized polysaccharides (**Figure 7C**) was compared with glycogen from rabbit liver (**Figure 7E**). As control, the amounts of free linear glucans were estimated by analyzing the APTS-labeled samples not incubated with commercial debranching enzyme (**Figures 7D and 7F**). In absence of significant amount of free glucan chains (**Figure 7D**), the *in vitro* synthesized polysaccharide harbors a typical CLD similar to animal glycogen with monomodal distribution and maltohexaose (DP6) as most abundant glucan chains. Altogether, these results confirm that GlgE activities display an *in vitro* function similar to that of glycogen synthase (GlgA) for initiating and elongating the growing glycogen particles.

### Expression of bifunctional TreS-Mak of *Estrella lausannensis*

To our knowledge, the characterization of the bifunctional TreS-Mak activity has not yet been reported in the literature. At variance to previous GlgE expression experiments, first transformation experiment in Rosetta^TM^ (DE3) *E.coli* strain did not yield colonies in spite of the absence of inducer (IPTG) (**S7A Figure**). We presumed that a basal transcription of *treS-mak* gene associated with a substantial intracellular amount of trehalose (estimated at 8.5 mM in *E.coli* cell spread on Luria-Broth agar medium [40]) lead to the synthesis of highly toxic maltose-1-phosphate. This encouraging result prompted us to perform expression of TreS-Mak protein in BL21-AI strain. The his-tagged TreS-Mak protein purified on nickel columns displays a molecular weight of 115 kDa on SDS-PAGE (**S7B Figure**) while in solution recombinant TreS-Mak formed a homodimer with an apparent molecular weight of 256 kDa as analyzed by sepharose 6 column chromatography (**S7C Figure**). This contrast with the hetero-octameric complex of 4TreS-4 Mak (≈490 kDa) observed in *Mycobacterium smegmatis* in which homotetramers of TreS forms a platform to recruit dimers of Mak *via* specific interaction domain [41, 42].

We first confirmed that the N-terminus TreS domain is functional by measuring the interconversion of trehalose into maltose. The amount of maltose was enzymatically quantified using the amyloglucosidase assay. Previous reports indicated that TreS activities are partially or completely inhibited with 10 mM of divalent cation while a concentration of 1 mM has positive effects. The effect of Mn^2+^ cation on the activity of TreS domain was inferred at 200 mM of trehalose. As depicted on **Figure 8A**, the activity of the TreS domain increases only slightly by 1.1-fold from 0 mM to 1 mM of Mn ^2+^ (0.37 µmol maltose. min^-1^.mg^-1^) whereas a noticeable decrease of TreS activity (0.24 µmol maltose. min^-1^.mg^-1^) is obtained at 10 mM of Mn ^2+^. As reported in the literature, the TreS activity is also associated with the release of glucose during the interconversion of trehalose into maltose. Because TreS activity is fused with the Mak domain in *E. lausannensis*, we tested the effect of a wide range of concentration of ATP concentration on the interconversion of trehalose (**Figure 8B**). Although no significant effect of ATP was observed on TreS activity at 1mM (0,43 µmol maltose. min^-1^.mg^-1^), TreS activity decreased by 0.6-fold at 3 mM-10 mM ATP (0.29µmol maltose. min^-1^.mg^-1^) and dropped by 2.8-fold when the ATP concentration reaches up to 20 mM (0,15 µmol maltose. min^-1^.mg^-1^). Finally, the apparent Km value for trehalose was determined at 42.3±2.7 mM in the presence of 1 mM MnCl_2_ and 0 mM ATP (**Figure 8C**). This is consistent with the apparent Km values for trehalose (50 to 100 mM) reported in the literature for TreS activity in various species [43].

**Figure 8:**
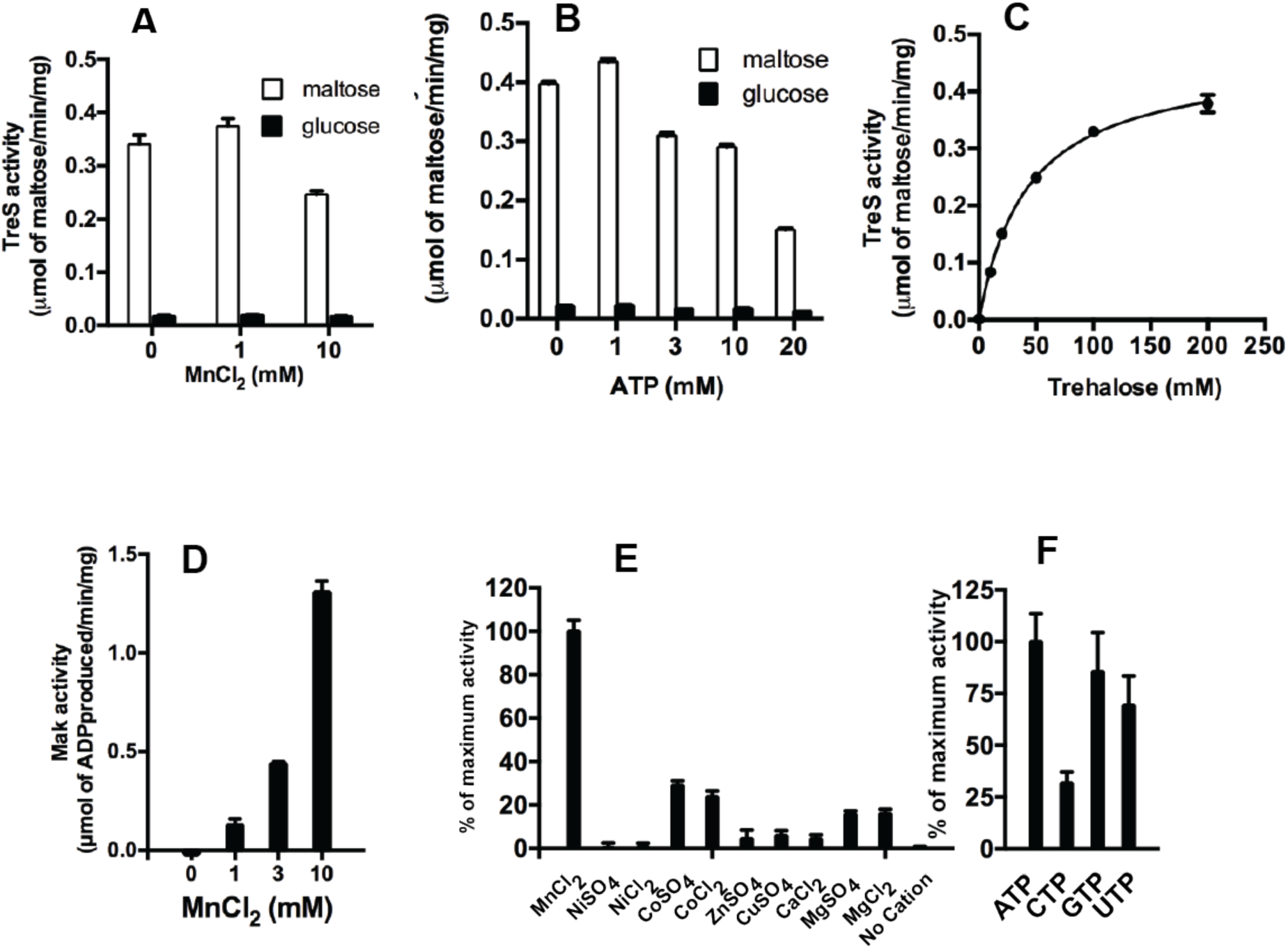
Biochemical properties of recombinant TreS-Mak of *Estrella lausannensis*. (**A**) The trehalose synthase domain (TreS) of bifunctional TreS-Mak activity was first conducted at 30°C pH 8 with 200 mM trehalose in presence of 0 mM, 1 mM and 10 mM of manganese chloride (MnCl_2_). The interconversion of trehalose into maltose and subsequent release of glucose were inferred by using amyloglucosidase assay method. The TreS activity is expressed as µmol of maltose/min/mg of protein. (**B**) The effect of nucleoside triphosphate on TreS activity was determined by measuring the interconversion of trehalose into maltose in the presence of increasing concentration of ATP (0 to 20 mM) and 200 mM trehalose. (**C**) The apparent Km value for trehalose was determined in the absence of ATP and 1 mM of MnCl_2_ by measuring the interconversion of increase concentration of trehalose (0 to 200 mM) into maltose. (**D**) Maltokinase activity domain was inferred by measuring the release of ADP during phosphorylation of maltose (20 mM) into M1P in presence of 0, 1, 3, 10 mM of MnCl_2_. The Mak activity is expressed as µmol of ADP released/min/mg of protein. (**E**) The effects of divalent cation Mn^2+^, Ni^2+^, Co^2+^, Zn ^2+^, Cu ^2+^, Ca^2+^, Mg^2+^ at 10 mM and (**F**) ATP, CTP, GTP and UTP nucleotides on Mak activity were determined and expressed as relative percentage of maximum activity.

We further focused on the activity of the maltokinase domain that catalyzes the phosphorylation of maltose in presence of ATP and releases M1P and ADP. The latter was monitored enzymatically *via* the pyruvate kinase assay in order to express the Mak activity domain as µmol of ADP released.min^-1^. mg^-1^ of protein. Our preliminary investigations indicated that imidazole stabilizes or is required for the maltokinase activity domain (**S7D Figure**). Hence all incubation experiments have been conducted in the presence of 125 mM of imidazole. The pH and temperature optima were respectively determined at 30°C and pH 7 (**S7E and S7F Figure**). Interestingly, the activity of the Mak domain is functional within a wide range of temperature that reflects probably the temperature of free-living amoebae or animal hosts. Kinase activities are reported for their requirement in divalent cation in order to stabilize the negatively charged phosphate groups of phosphate donors such as ATP. Therefore, TreS-Mak activity was inferred in the presence of various divalent cations (**Figure 8E**). As expected, the recombinant TreS-Mak was strictly dependent on divalent cations, in particular, with a noticeable stimulatory effect of Mn ^2+^ (**Figure 8D and 8E**). Others tested divalent cations, like Co^2+^, Mg^2+^, Fe^2+^, Ca^2+^, Cu^2+^ activated the Mak activity as well, but to a lower extent, while no effect was observed in presence of Ni^2+^. Interestingly, at variance to Mak activity of *Mycobacterium bovis*, which prefers Mg ^2+^, the catalytic site of the Mak activity domain of TreS-Mak binds preferentially Mn ^2+^ over Mg ^2+^ [44], which is consistent with a distinct evolutionary history as depicted on **figure 2B**. Then, nucleotides, ATP, CTP, GTP and UTP were tested as phosphate donors by measuring the amount of M1P released (**Figure 8F**). The data expressed in percentage of activity show that ATP (100%), GTP (85%), UTP (70%) and to a lower extent CTP (31%) are efficient phosphate donors. Altogether, we demonstrated that TreS and Mak domains are functional in the fused protein TreS-Mak of *E. lausannensis*. The reversible interconversion of trehalose combined with an intracellular trehalose concentration probably below 42 mM (the intracellular trehalose concentration was estimated at 40±10 mM inside one cell of *E. coli* strain overexpressing OtsA/OtsB [40]) suggest that irreversible phosphorylation of maltose drives the synthesis of M1P.

### *Estrella lausannensis* and *Waddlia chondrophila* accumulate glycogen particles within the cytosol of EB via GlgE-pathway

Since incubation experiments have shown that branched polysaccharide can be synthesized in the presence of maltose-1-phosphate and both GlgE and GlgB activities, this prompted us to examine the presence of glycogen particles in thin section of *E. lausannensis* and *W.chondrophila* by transmission electron microscopy. After twenty four hours post infection, thin sections of *Acanthamoeba castellanii* infected with both *Chlamydiales* (**Figures 9A and 9C**) and purified elementary bodies (**Figures 9B and 9D**) were subjected to specific glycogen staining based on the periodic acid method, which is considered to be one of the most reliable and specific methods for staining glycogen [45]. Glycogen particles appear as electron-dense particles (white head arrows) in the cytosol of elementary bodies of *E. lausannensis* and *W. chondrophila*. Interestingly, because *Waddlia chondrophila* infects animal cells, which do not synthesize trehalose, this suggests that trehalose must be synthesized by the bacteria itself [46]. Based on the five different trehalose pathways described in prokaryotes (for review [47]), we found that trehalose biosynthesis is limited to so-called “environmental *Chlamydiae*” and is not present in the *Chlamydiaceae* family. Among chlamydial strains with GlgE pathway, *P. phocaeensis* and *P. neagleriophila* synthesize trehalose through TreY-TreZ pathway while OtsA-OtsB pathway was found in both *E. lausannensis* and *W. chondrophila.* Importantly, *otsA* and *otsB* genes encode for trehalose-6-phosphate synthase and trehalose-6-phosphate phosphatase, respectively. OtsA activity condenses glucose-1-phosphate and UDP-glucose into trehalose-6-phosphate. However, BLAST search did not evidence the classical *galU* gene encoding UDP-glucose pyrophosphorylase, which synthesizes UDP-glucose from glucose-1-phosphate and UTP in both *E. lausannensis* and *W. chondrophila*, but rather a non-GalU type UDP-glucose pyrophosphorylase homolog to UGP3 of plants. This chloroplastic UDP-glucose pyrophosphorylase activity, dedicated to sulfolipid biosynthesis belongs to a set of 50 to 90 chlamydial genes identified, as lateral gene transfer, in the genomes of Archaeplastida [48]. Based on this work and taking into account the current genome analysis, we propose that glycogen metabolism pathway in *W. chondrophila* and *E. lausannensis* occur as depicted on **Figure 10**.

**Figure 9:**
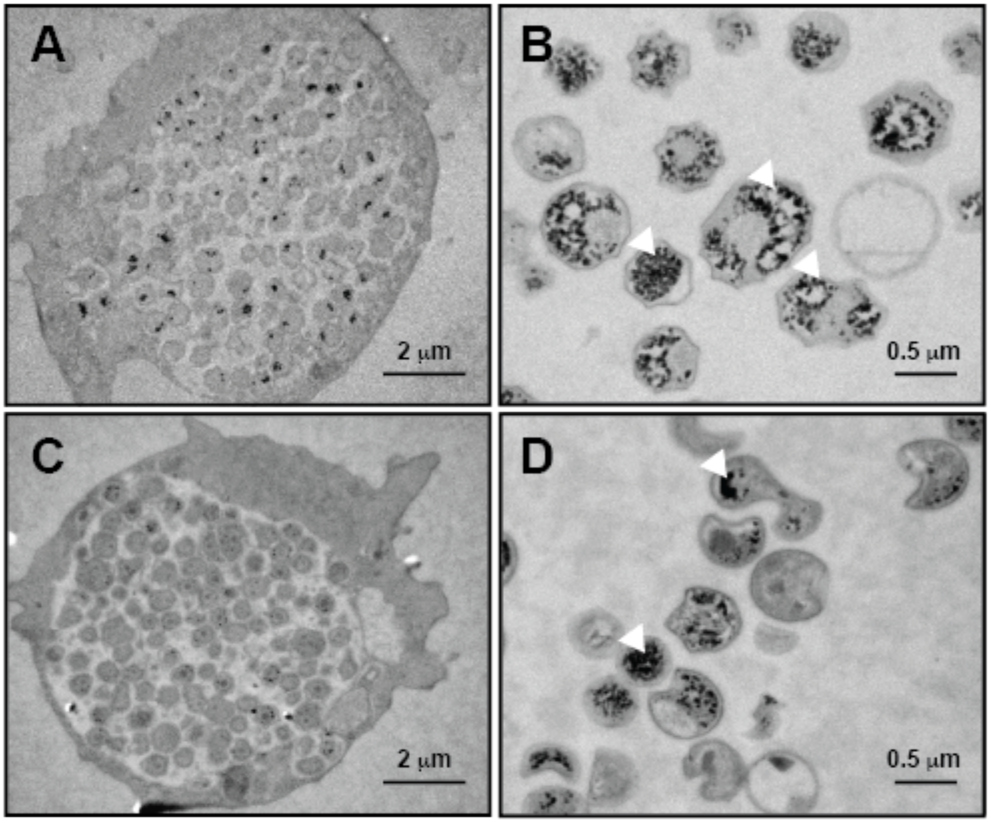
Glycogen accumulation in *Estrella lausannensis*. (A, B) and *Waddlia chondrophila* (C, D). Glycogen particles (white head arrows) in *E.lausannensis* and *W. chondrophila* were observed by TEM after periodic acid thiocarbohydrazide-silver proteinate staining of ultrathin sections of 24h post infected *A. castellanii* with *E. lausannensis* (A) and *W. chondrophila* (C) or purified bacteria (B, D).

**Figure 10:**
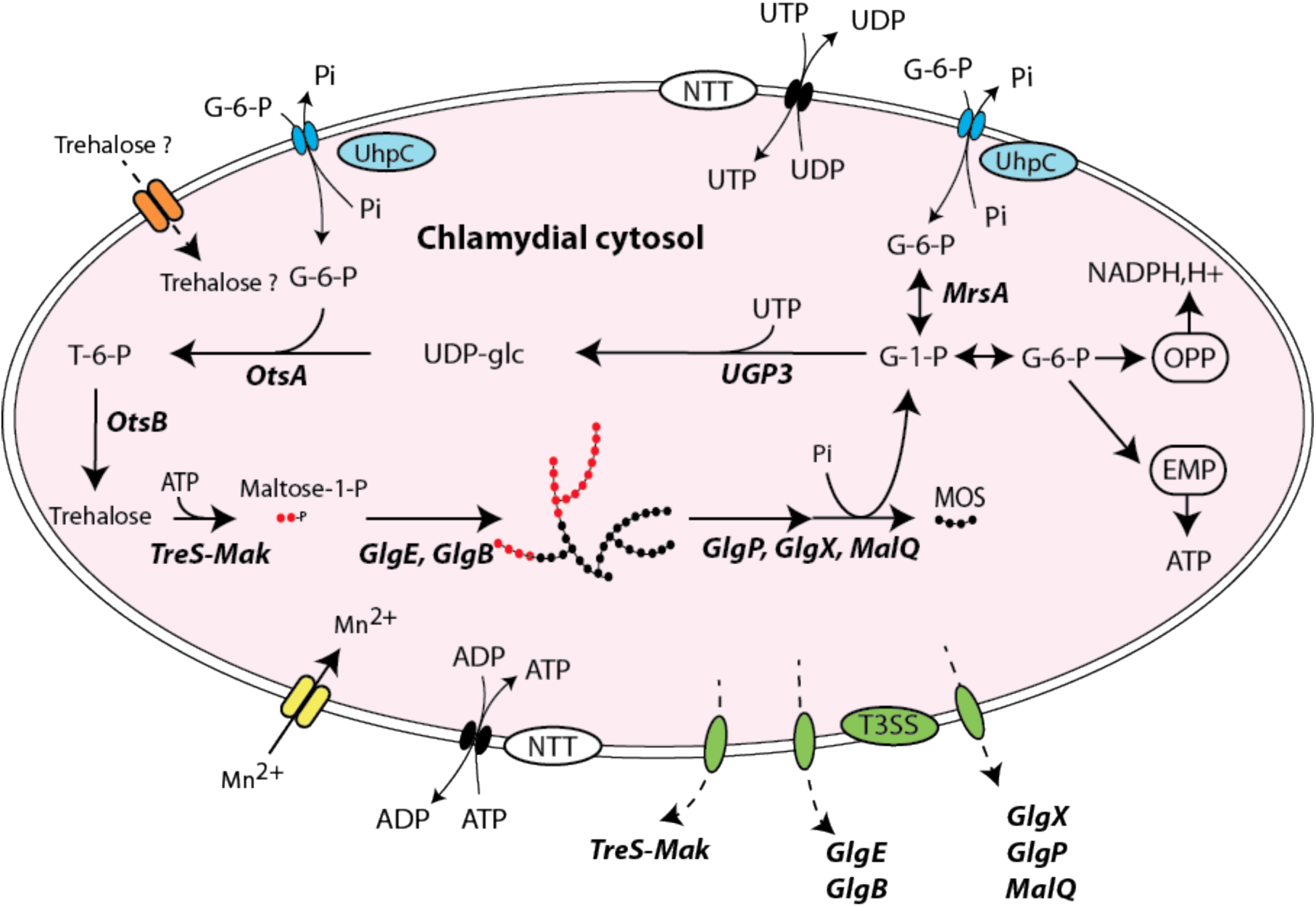
Glycogen metabolism network in *Waddliaceae* and *Criblamydiaceae* families. Glucose-6-phosphate (G-6-P) and UTP/ATP are transported in the cytosol via Uhpc and NTT translocators. The first committed step consists into the isomerization of G-6-P into glucose-1-phosphate (G-1-P) catalyzed by glucose-6-phosphate isomerase activity (***MrsA***). UDP-glucose pyrophosphorylase (***UGP3***) synthesizes UDP-glucose from G-1-P and UTP. Both trehalose-6-phosphate synthase (**OtsA**) and trehalose-6-phosphate phosphatase (**OtsB**) convert nucleotide-sugar and G-6-P into trehalose. The bifunctional TreS-Mak activity supplies the maltosyl transferase activity (**GlgE**) in maltose-1-phosphate (M1P). *De novo* glucan initiation and elongation properties of GlgE and branching enzyme activity (**GlgB**) allow the appearance of α-polysaccharide (*i.e.* glycogen) made of α-1,4 and α-1,6 linkages. The synergic action of glycogen phosphorylase (**GlgP**), debranching enzyme (**GlgX**) and a-1,4 glucanotransferase (**MalQ**) depolymerize glycogen into G-1-P and short malto-oligosaccharides (MOS). The former fuels both oxidative pentose phosphate (OPP) and Embden-Meyerhof-Parnas (EMP) pathways that supply the extracellular forms (elementary bodies) in reduced power (NADPH,H+) and ATP, respectively. Divalent cations Mn^2+^ required for TreS-Mak activity are probably imported *via* ABC transporter composed of MntA, MntB and MntC 3,sub-units identified in chlamydial genomes. *Waddliaceae* and *Criblamydiaceae* may manipulate the carbon pool of the host by uptaking trehalose through a putative disaccharide transporter (orange/dash arrow) or by secreting glycogen-metabolizing enzymes through type three-secretion system (green/dash arrow).

## Discussion

The present study examined the glycogen metabolism pathway in *Chlamydiae* phylum. Unlike other obligatory intracellular bacteria, *Chlamydiae* have been documented to retain their capacity to synthesize and degrade the storage polysaccharide with the notable exception of the *Criblamydiaceae* and *Waddliaceae* families, for which the key enzyme of glycogen biosynthesis pathway, ADP-glucose pyrophosphorylase activity was reported missing [11, 49]. All mutants deficient in GlgC activity are associated with glycogen-less phenotypes and so far no homologous gene encoding for a GlgC-like activity has been established among prokaryotes [50, 51]. To our knowledge, only two cases have been documented for which GlgC activity has been bypassed in the classical GlgC-pathway. The ruminal bacterium *Prevotella bryantii* that does not encode an ADP-glucose pyrophosphorylase (*glgC*) gene has replaced the endogenous *glgA* gene with an eukaryotic UDP-glucose-dependent glycogen synthase [52, 53]. The second case reported concerns the GlgA activity of *Chlamydia trachomatis* which has evolved to polymerize either UDP-glucose from the host or ADP-glucose produced by GlgC activity into glucose chains [54]. In order to get some insight in *Criblamydiaceae* and *Waddliaceae* families, a survey of glycogen metabolizing enzymes involved in the classical GlgC-pathway and in the recently described GlgE-pathway was carried out over 47 chlamydial species representing the diversity of chlamydiae phylum. As expected, we found that a complete GlgC-pathway in most chlamydial families and the most astonishing finding was the occurrence of GlgE-pathway in three phylogenetically related *Parachlamydiaceae*, *Waddliaceae* and *Criblamydiaceae* families. Our genomic analysis also pinpointed a systematic lack of *glgC* gene in 6 draft genomes of candidatus *Enkichlamydia* sp. Those genomes are derived from metagenomic studies and were estimated to be 71 to 97% complete. If we assume the loss of *glgC*, the characterization of glycogen synthase with respect to nucleotide sugars should shed light on the glycogen pathway, as a result provide another example of the bypassing of GlgC activity in the classical GlgC-pathway. In addition to the occurrence of GlgE-pathway, a detailed genomic analysis of *Waddliaceae* and *Criblamydiaceae* families has revealed a large rearrangement of *glg* genes of GlgC-pathway, which had led to the loss of *glgP* gene and a fusion of *glgA* and *glgB* genes. This fusion appears exceptional in all three domains of life and no other such examples have been reported. In *E. lausannensis* (*Criblamydiaceae* fam.), the fusion of *glgA-glgB* genes is associated with a non-sense mutation resulting in premature stop codon in the open reading frame of the GlgA domain precluding the presence of the fused branching enzyme. We have shown that the glycogen synthase domain of chimeric GlgA-GlgB of *W. chondrophila* was active and remained ADP-glucose dependent while its branching enzyme domain already appears to be non functional due to the presence of GlgA domain at the N-terminal extremity that prevents the branching enzyme activity. In line with these observations, at variance with *Parachlamydiaceae* which have only maintained the *glgB* gene of GlgC pathway, both *Waddliaceae* and *Criblamydiaceae* have conserved *glgB2* gene in GlgE operon, thereby, further suggests that the glycogen branching activity domain is indeed defective or impaired in all GlgA-GlgB fusions. Overall, this study clearly implies that GlgC-pathway does not operate in both the *Waddliaceae* and *Criblamydiaceae* families. In addition it appears possible that the genes required for the presence of a functional GlgC-pathway are at different stages of disappearance from these genomes, as suggested by the non-sense mutation in glgA-glgB gene of *E. lausannensis* [55].

We further investigated the GlgE glycogen biosynthesis pathway in *Chlamydiales*. A series of biochemical characterizations have shown that GlgE activities are capable of transferring maltosyl residue of maltose-1-phosphate onto linear chain of glucose. More remarkably, GlgE activities fulfill the priming function of glycogen biosynthesis as described for GlgA activity in GlgC-pathway [56]. We have shown that the GlgE activities switch between the processive or distributive modes of polymerization depending on the initial presence of glucan chains. Thus the “processive mode” of GlgE activity yields long glucan chains (DP>32) and is favored in their absence or in the presence of short glucan primers (DP<4). This “processive mode” of GlgE activity fills up the critical function of initiating long glucan chains that will be taken in charge by the branching enzyme in order to initiate the formation of glycogen particles. *In vitro* incubation experiments performed in the presence of M1P and/or branching enzyme activity further confirmed that GlgE activity is by itself sufficient for synthesizing *de novo* a branched polysaccharide with high molecular weight. At variance with mycobacteria and *Streptomycetes*, trehalose synthase (TreS) and maltokinase (Mak) activities of *Chlamydiales* form a bifunctional enzyme composed of TreS and Mak domains at the N- and C-terminus, respectively, which has never been reported to our knowledge. The fused TreS-Mak activity is functional and mediates the trehalose conversion into maltose and the phosphorylation of maltose into maltose-1-phosphate in the presence of ATP, GTP or UTP as phosphate donors. In contrast to mycobacteria, the maltose kinase domain requires preferentially manganese rather magnesium as divalent cation [44].

The fact that the occurrence of GlgE-pathway is limited to a few chlamydial families has led us to wonder about the origin of this operon. Our phylogeny analyses suggest that GlgE operons identified in chlamydia species share a common origin but are only distantly related to the GlgE operon from Actinobacteria (i.e Mycobacteria). We could not determine whether the presence of the GlgE pathway predated the diversification of chlamydiae or whether the operon was acquired by lateral gene transfer by the common ancestor of the *Criblamydiaceae*, *Waddliaceae* and *Parachlamydiaceae* from another member of PVC superphylum. One fair inference is that the genome of common ancestor of *Waddliaceae, Criblamydiaceae* and *Parachlamydiaceae* families encoded both GlgC- and GlgE-pathways. The loss of both glgP and glgC and fusion of glgA and glgB occurred before the emergence of *Waddliaceae* and *Criblamydiaceae* and may involve one single deletion event if we presume a *glA/glgC/glgP/glgB* gene arrangement in the common ancestor. While GlgE pathway was maintained in *Waddliaceae* and *Criblamydiaceae* due to the mandatory function of glycogen in *Chlamydiales*, most of members of *Parachlamydiaceae* retained only the GlgC-pathway except for two *Protochlamydia* species. The redundancy of glycogen metabolism pathway in *P. naegleriophila* species is quite surprising and goes against the general rule of genome optimization of intracellular obligatory bacteria. It is worthy to note that *P. naegleriophila* species was originally isolated from a protist *Naegleria* sp. *N. fowleri,* the etiological agent of deadly amoebic encephalitis in humans, stores carbon exclusively in the form of trehalose and is completely defective for glycogen gene network [57]. Therefore, it is tempting to hypothesize that *P. neagleriophila* use the retained the GlgE pathway to effectively mine the trehalose source of its host either by uptaking trehalose from its host through a putative disaccharide transporter or by secreting via type three secretion system enzymes of GlgE pathway. Our preliminary experiments based on heterologous secretion assay in *Shigella flexneri* suggested that GlgE and TreS-Mak could be secreted by the type three-secretion system (**S8 Figure)**. Like *Chlamydiaceae*, the secretion of chlamydial glycogen metabolism pathway may be a strategy for manipulating the carbon pool of the host [54]. As reported for *P. amoebophila* with respect to D-glucose [49], the utpake of host’s trehalose provides an important advantage in terms of energy costs. In comparison with GlgC-pathway, one molecule of ATP is required to incorporate two glucose residues onto growing polysaccharide; at the scale of one glycogen particle synthesis this may represent a significant amount of ATP saving. The uptake of radiolabeled trehalose by host-free elementary bodies may or not support this hypothesis.

The preservation of glycogen metabolism pathway through the bottleneck of genome reduction process sheds light on a pivotal function of glycogen that has been hitherto underestimated within Chlamydiae. As a result, the question arises as to why chlamydiae have maintained glycogen metabolism pathway, making them unique among obligate intracellular bacteria. It is worthy to note that most of obligate intracellular bacteria *Anaplasma* spp., *Ehrlichia* spp., *Wolbachia* spp., *Rickettsia* spp. do not experience environmental stresses like Chlamydiae and *Coxellia burnetii* [58]. They thrive in nutrient rich environments either in animal or insect hosts. Losses of metabolic functions such as carbon storage metabolism in obligate intracellular bacteria are balanced by the expression of a wide variety of transporters for the uptake metabolites from the host. Except for ultra-resistant spore-like forms of *C. burnetti* named small cell variants, over the last decade, our perception of EB has switched from an inert spore-like form to metabolic active form capable of transcription and translation activities [49, 59]. The combination of different “omic” approaches performed on purified RB and EB of *C. trachomatis* and *P. amoebophila* have shown that genes involved in glycogen and energy metabolism pathways are upregulated in the late stage of development [4] [60][61] and most remarkably, the uptake of glucose and glucose-6-phosphate by EBs of *P. amoebophila* and *C. trachomatis* improves significantly the period of infectivity [49, 62]. Accordingly, it seems reasonable to argue that the primary function of cytosolic glycogen in EBs is to fuel metabolic processes (i.e glycolysis, pentose phosphate) when EBs are facing up the poor nutrient environment (**figure 10**). Future investigations should provide new opportunity to delineate the function of glycogen in chlamydiae especially with the development of forward genetic approaches [63, 64]. Finally the use of GlgE inhibitors initially designed against mycobacterial infections and to some extent the use of inhibitors of chlamydial glycogen metabolizing enzymes might define new attractive drugs to treat *W. chondrophila,* since this *Chlamydia-*related bacteria has been increasingly recognized as a human pathogen [65, 66].

## Acknowledgements

The authors are very grateful to Dr Nicolas Szydlowski for providing access to the capillary electrophoresis. We also thank Dr Agathe Subtil from Pasteur Institute (Paris) for providing *Shigella* strains and antibodies as well as the Plateforme d’Analyse des Glycoconjugues (PAGes, http://plateforme-pages.univ-lille1.fr/) for providing access to the instrumental facilities for carbohydrate analysis. This work was supported by the CNRS, the Université de Lille CNRS, and the ANR grants “Expendo” (ANR-14-CE11-0024).

---

**S1 figure:**
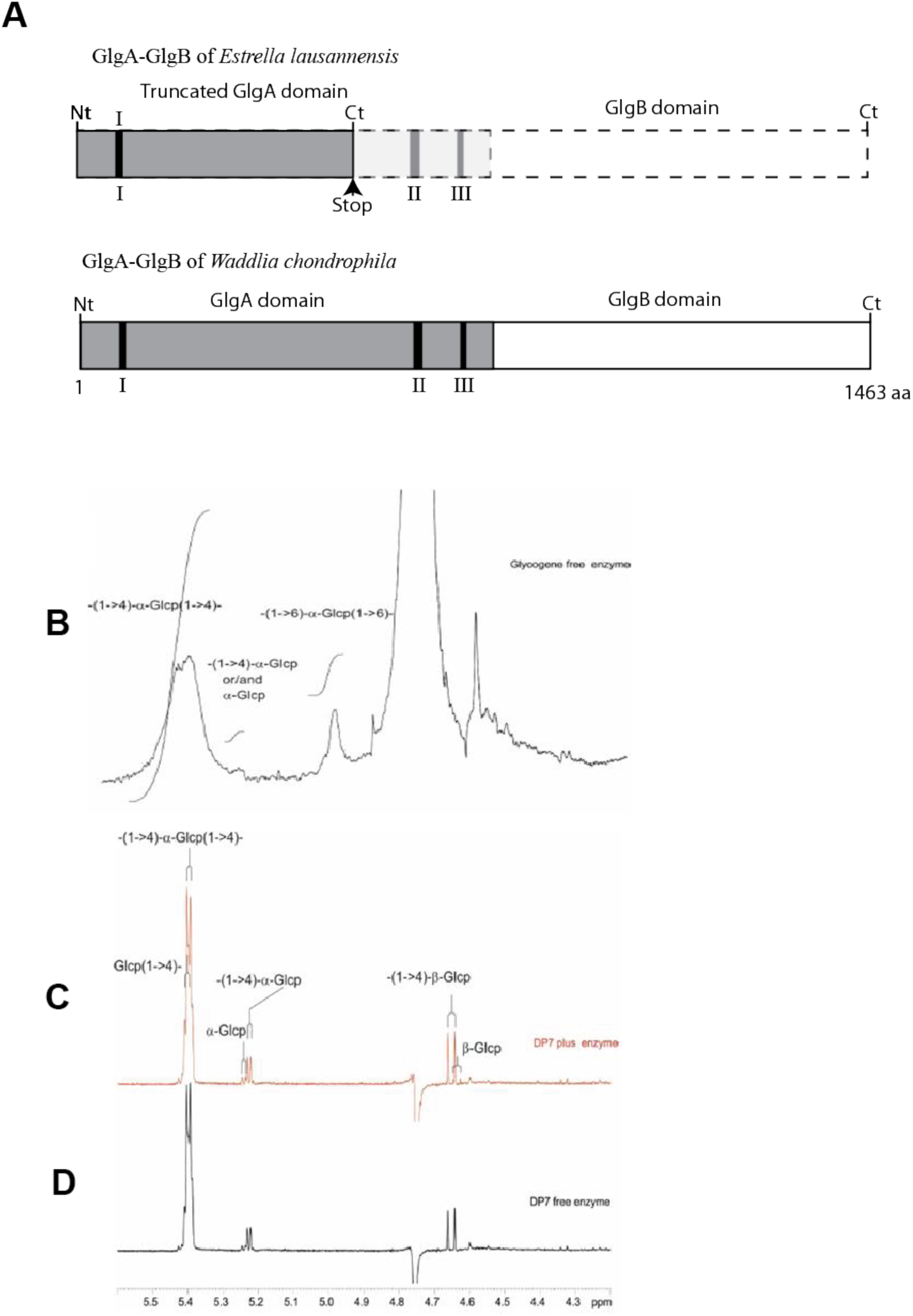
(**A**) Domain organization of fused protein GlgA-GlgB of *E. lausannensis* and *W. chondrophila*. Glycogen synthase domain (gray box) and branching enzyme domain (white box) are respectively located at the N-(Nt) and C-termini (Ct) respectively. The insertion of one-nucleotide in *E. lausannensis* sequence results in a frame shift and the appearance of truncated GlgA-GlgB protein. Regions I, II and III represent highly conserved sequences in the glycogen synthase GT5 family that includes amino acid residues involved in the catalytic site and nucleotide binding sites. Proton-NMR analyses of glycogen from rabbit liver (**B**) and maltoheptaose (10 mg.mL^-1^) + ADP-glucose (3 mM) incubated overnight at 30°C in the presence (**C**) or in the absence (**D**) of recombinant GlgA-GlgB fusion enzyme of *Waddlia Chondrophila.* After incubation, enzymatic reactions were boiled and purified through anion and cation exchange resins (DOWEX 1 × 8 and DOWEX 50 W X 8). Protons involved in α-1,4 linkages or α-1,6 linkages resonate at 5.4 and 4.95 ppm, respectively. Protons in α and β position on C1 (reducing end) generate signals at 5.23 and 4.65 ppm. The absence of signal at 4.95 ppm suggests that either signal corresponding to α-1,6 linkages is below threshold of detection (<1%) or GlgB domain is not active in the GlgA-GlgB of WC.

**S2 figure:**
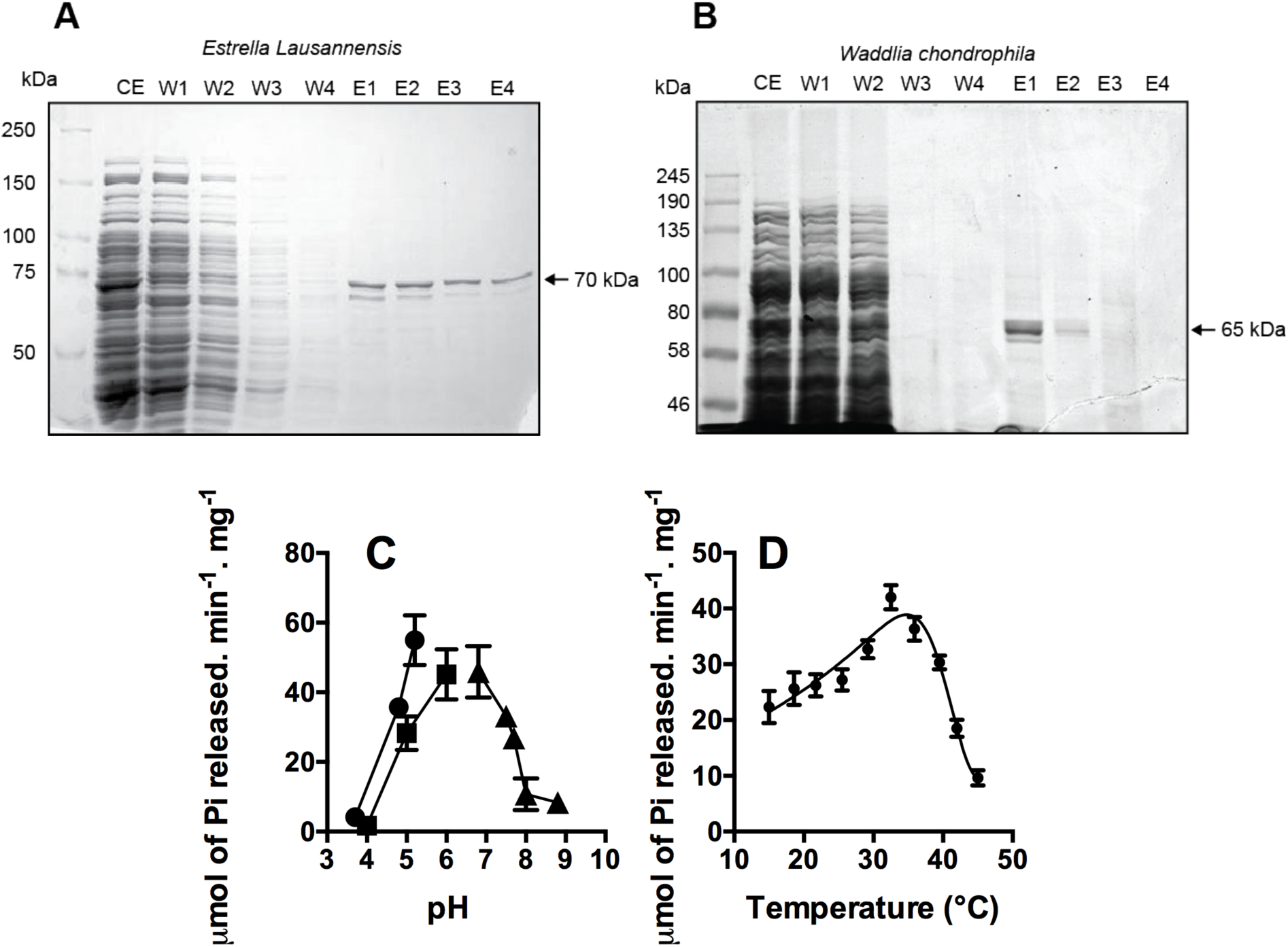
SDS-PAGE analyses of recombinant GlgE after affinity column purification and determination of optima pH and temperature of GlgE-EL. His-tag GlgE of *E. lausannensis*. (A) and *W. chondrophila* (B) were expressed in Rosetta^TM^ *E. coli* strain. After induction at mid exponential growth with IPTG for GlgE-EL and culture in auto-inductible medium for GlgE-WC, the overnight cultures were harvested by centrifugation. Cell pellets were suspended in loading buffer containing 25 mM TRIS/acetate pH 7.5 and then subjected to sonication. After centrifugation, crude extract (CE) was incubated with nickel affinity column at 4°C for one hour. Total proteins in both CE and affinity purification fractions; flow-through (FT), washing steps (W1 to W4) and elution (E1 to E4) fractions were separated on SDS-PAGE 7.5%. Based on standard molecular weights, the apparent molecular weights of GlgE were estimated at 76 kDa and 72 kDa for *E. lausannensis* and *W. chondrophila*, respectively. The optima of pH (C) and temperature (D) of GlgE of *E. lausannensis* were determined by measuring the amount of orthophosphate released after the transfer of maltosyl moieties of M1P onto the non-reducing end of glucan chains. The optimum pH determination was carried out at 30°C in sodium acetate ((circle), pH 3.7; 4.8; 5.2), sodium citrate ((square), pH 4; 5; 6) and TRIS/HCl ((triangle) pH 6.8; 7.5; 7.7; 8; 8.8) buffers with a final concentration of 25mM. The optimum pH determination was carried out in the presence of 25 mM TRIS/HCl pH 6.8.

**S3 figure:**
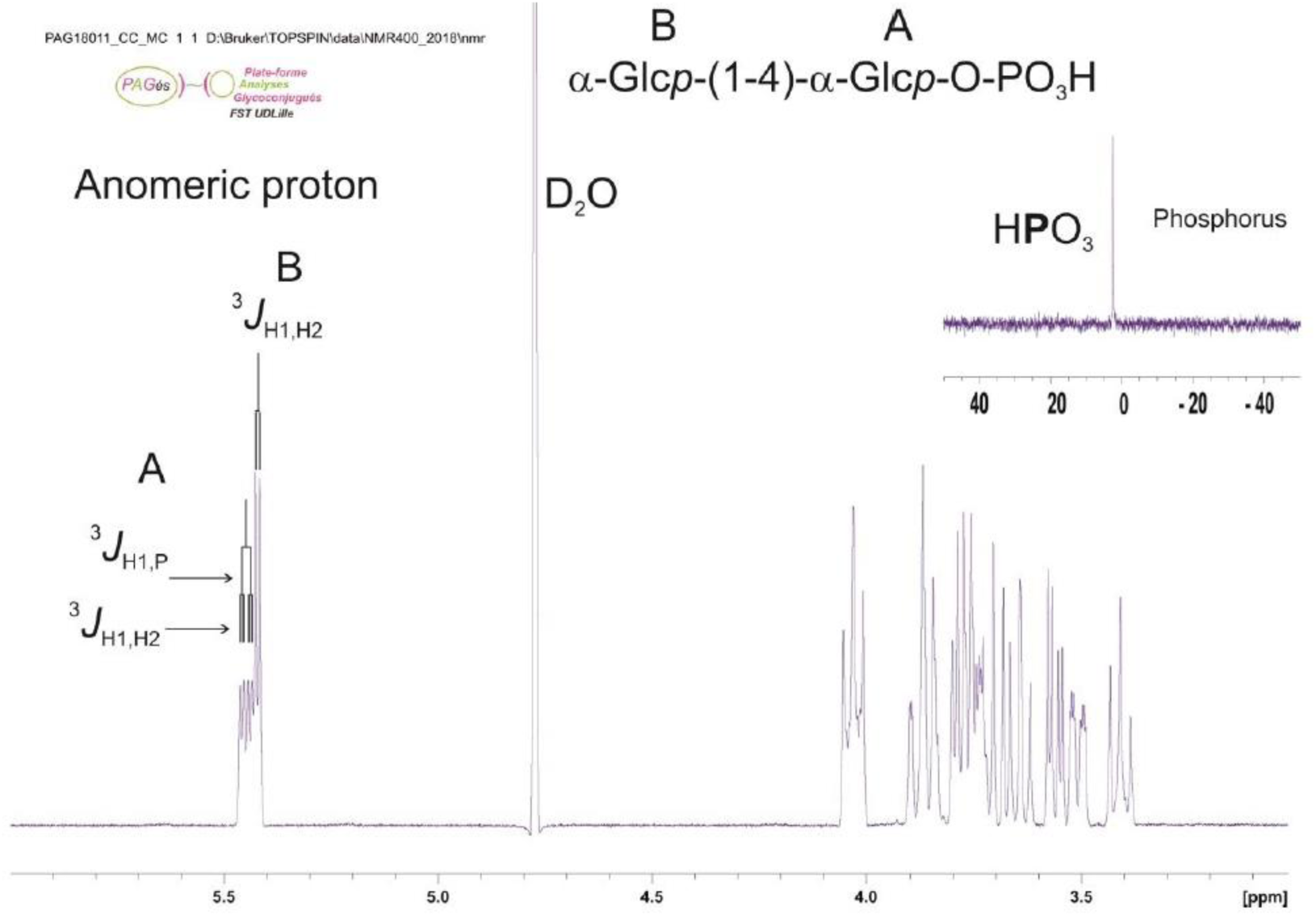
Proton- and phosphate-NMR analyses. Reaction product purified following the incubation of Waddlia chondrophila GlgE with glycogen and orthophosphate. Complete 1D-^1^H-NMR spectrum of maltoside-1-phosphate. α-anomer configuration of both glucosyl residues were characterized by their typical homonuclear vicinal coupling constants (^3^*J*_H1A,H2A_ and ^3^*J*_H1B,H2B_) with values of 3.5 Hz and 3.8 Hz respectively. A supplementary coupling constant was observed for α-anomeric proton of residue A as shown the presence of the characteristic doublet of doublet at 5.47 ppm. This supplementary coupling constant is due to the heteronuclear vicinal correlation (^3^*J*_H1A,P_) between anomeric proton of residue A and phosphorus atom of a phosphate group, indicating that phosphate group was undoubtedly *O*-linked on the first carbon of the terminal reducing glucosyl unit A. The value of this ^3^*J*_H1A,P_ was measured to 7.1Hz.

**S4 figure:**
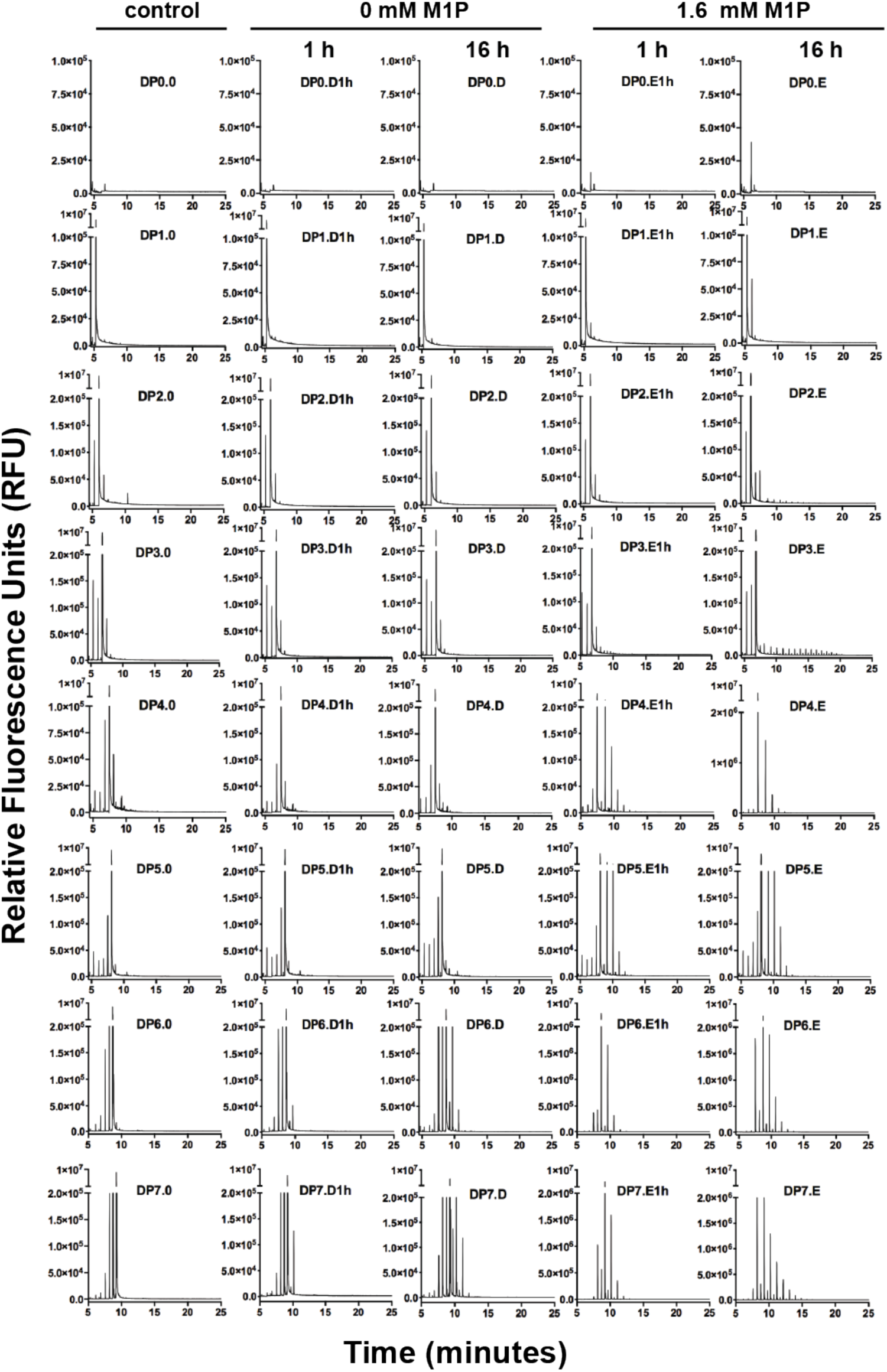
FACE analysis of activity GlgE of *Estrella Lausannensis.* Recombinant GlgE (3.5 nmol Pi.min^-1^) was incubated 1 hour and 16 hours at 30°C with 5 mM of malto-oligosaccharides composed of 0 to 7 glucose moieties (degree of polymerization: DP) and 0 mM or 1.6 mM of maltose-1-phosphate (M1P). After incubation, enzymatic reactions are stopped 5 min at 95°C. Malto-oligosaccharides are labeled with APTS and then separated according to their DP using capillary electrophoresis. Fluorescence is monitored as relative fluorescence units (RFU). As control, heat denatured GlgE activity were incubated 16 hours at 30°C with M1P and malto-oligosaccharides.

**S5 figure:**
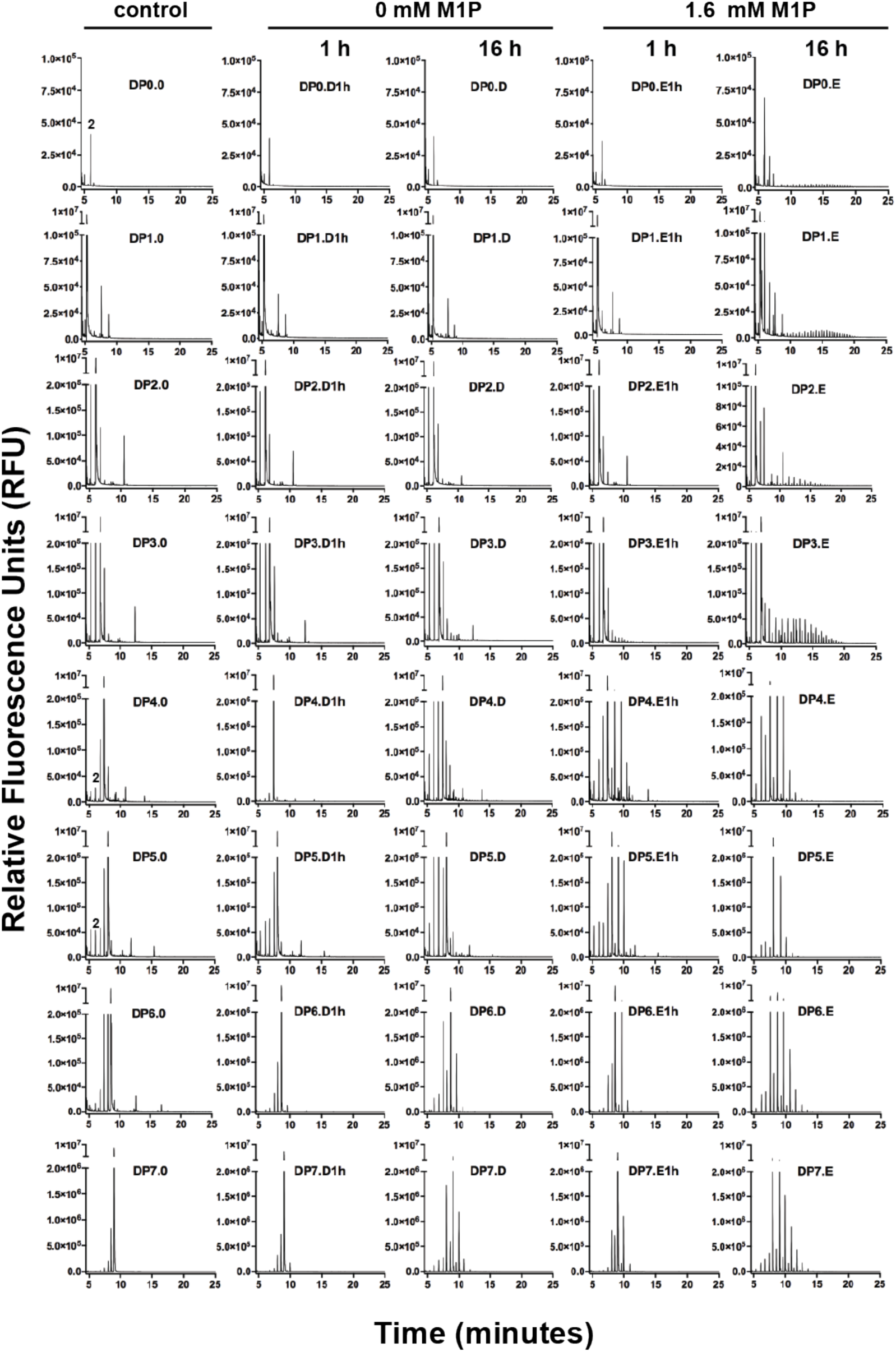
FACE analysis of activity GlgE of *Waddlia chondrophila.* Recombinant GlgE (1.38 nmol Pi.min^-1^) was incubated 1 hour and 16 hours at 30°C with 5 mM of malto-oligosaccharides composed of 0 to 7 glucose moieties (degree of polymerization: DP) and 0 mM or 1.6 mM of maltose-1-phosphate (M1P). After incubation, enzymatic reactions are stopped 5 min at 95°C. Malto-oligosaccharides are labeled with APTS and then separated according to their DP using capillary electrophoresis. Fluorescence is monitored as relative fluorescence units (RFU). As control, heat denatured GlgE activity were incubated 16 hours at 30°C with M1P and malto-oligosaccharides.

**S6 Figure:**
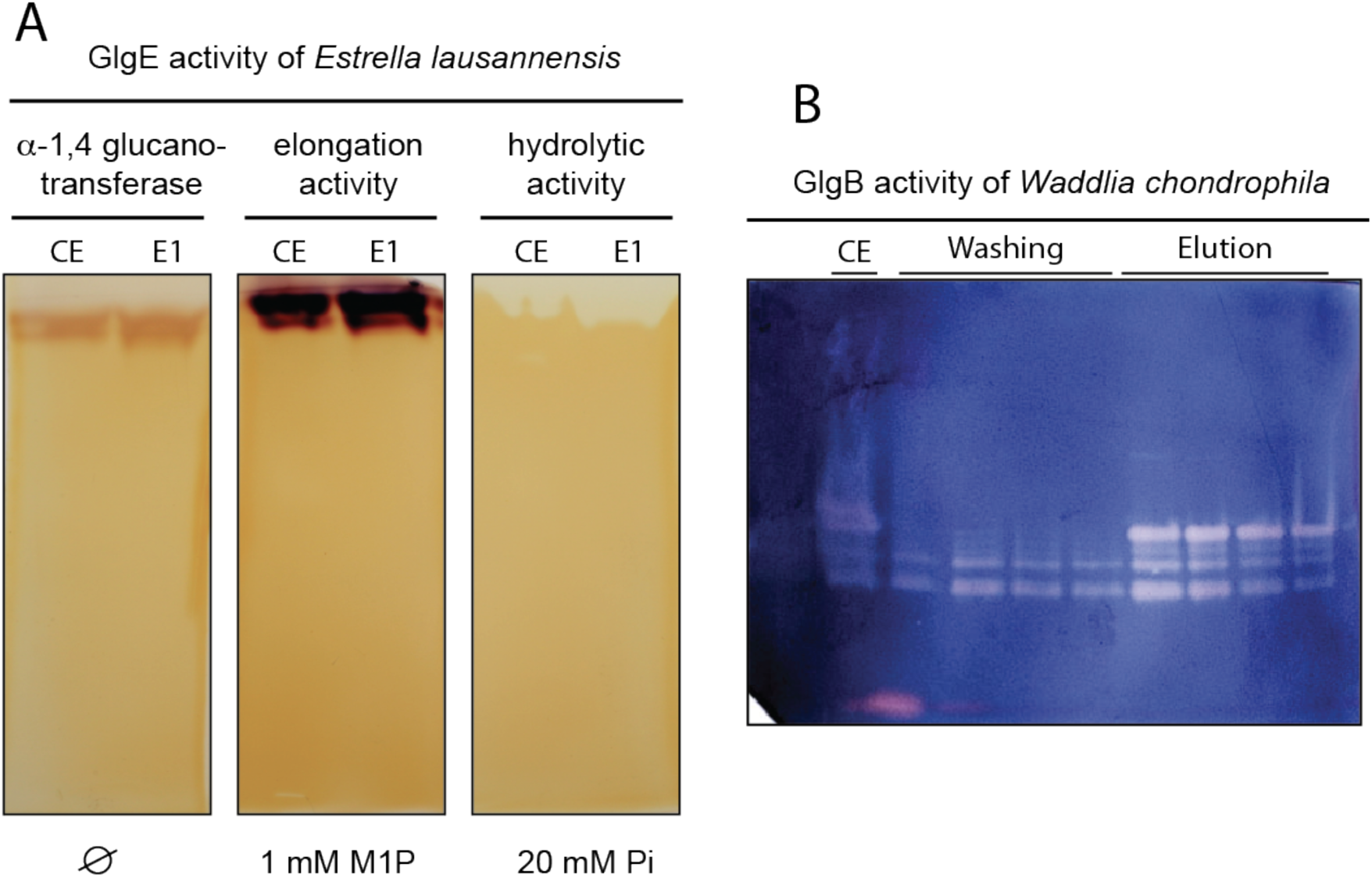
(A) Native-PAGE containing glycogen reveals α-1,4 Glucanotransferase, elongation and hydrolytic activities of *E. lausannensis* GlgE. *E. coli* crude extract (CE) expressing GlgE-EL and purified GlgE-EL fraction (E1) were loaded onto native-PAGE containing 0.3% (w/v) of glycogen from bovine liver. Gel runs in ice pocket during 1h30 at 15 mA constant in TRIS/glycine buffer pH 8.8. After electrophoresis, native-gel was cut in three pieces and incubated overnight at room temperature with 10 mL 25 mM TRIS/acetate buffer pH 7.5 (Ø), 10 mL 25 mM TRIS/acetate buffer pH 7.5 and 1 mM of maltose-1-phosphate (M1P), 10 mL 25 mM TRIS/acetate buffer pH7.5 and 20 mM orthophosphate (Pi). Soaking native gel in iodine solution evidences GlgE activity. α-1,4 Glucanotransferase activity is visualized as brownish activity bands due to maltosyl reaction transfers catalyzed by GlgE-EL on the external glucan chains of glycogen particles. In presence of 1 mM M1P, the elongation activity is favored and consists in the maltosyl moieties transfer reactions of M1P onto non-reducing ends of external glucan chains of glycogen. The increase of long glucan chains leads to a strong iodine-glucan interaction observed as a dark activity band. At contrary, the hydrolytic reaction is conducted in the presence of 20 mM of Pi. GlgE-EL releases M1P from the non-reducing ends of external glucan chains of glycogen and α-1,6 linkages or branching points prevent the complete hydrolysis of glycogen particles. Nevertheless, the resulting branched glucans escape from polyacrylamide gel leading to clear activity band in orange background. (B) Purification of branching enzyme (GlgB) activity of *Waddlia chondrophila*. The plasmid expression pET15b-GlgB-WC was transferred in ΔglgB Rosetta^TM^ *E. coli* strain impaired in endogenous branching enzyme. After induction, crude extract (CE) was incubated for one hour with his-agarose beads at 4°C. Unbound proteins were eluted with 50 mM sodium acetate, 300 mM NaCl and 60 mM imidazole pH 7. After four washing steps, His-GlgB were eluted with 50 mM sodium acetate, 300 mM NaCl and 250 mM imidazole pH 7. Proteins in the flow through and elution fractions were separated on native-PAGE (7.5%) at 4°C (120 V, 15 mA). After electrophoresis, proteins were electrotransfered against a native-PAGE containing 0.3% (w/v) of potato starch using Trans-Blot® Turbo™ transfer system (Bio-Rad). Native-PAGE was then incubated overnight in 25 mM TRIS/acetate buffer pH 7.5 at room temperature. Branching enzyme activity is revealed as pink bands in blue background after soaking the gel in iodine solution (KI/I_2_).

**S7 Figure:**
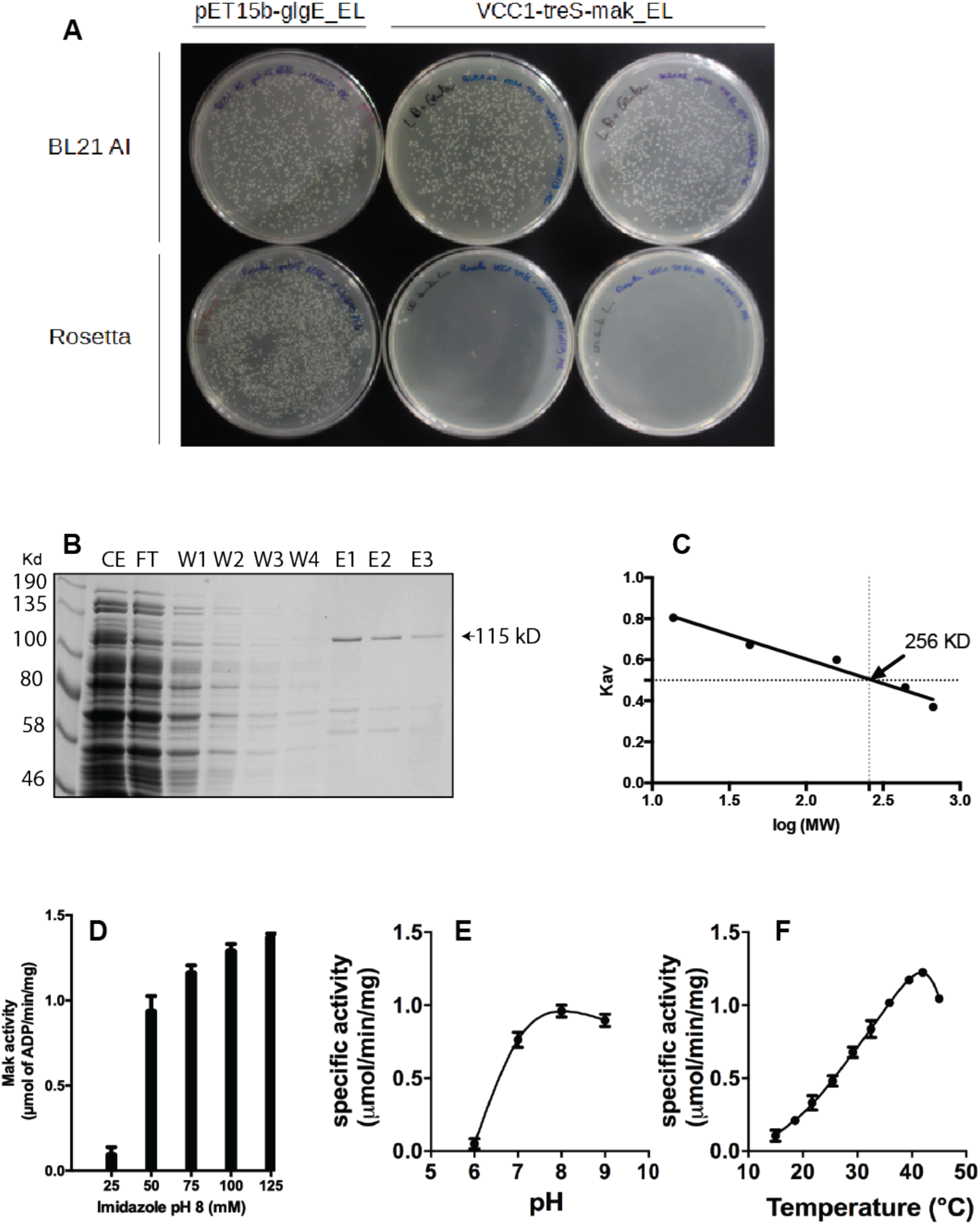
(**A**) Toxicity of TreS-Mak activity of *E. lausannensis*. Two chemically competent *E. coli* cells Rosetta^™^ and BL21-AI^™^ were transformed with 0.15 µg of expression plasmids VCC1-TreS-Mak-EL (7.2 kpb) as well as with 0.15 µg of pET15-GlgE-EL (7.9 kpb) used as a control. Following the transformation, cells were spread onto Luria Broth medium containing appropriate antibiotic. Despite the lack of inductor (IPTG), no Rosetta^TM^ colonies were observed after 16 hours at 37°C for two independent constructions of VCC1-TreS-Mak#1 and #2 while a close number of BL21-AI colonies are visualized with pET-15-GlgE and VCC1-TreS-Mak plasmids. The leaky transcriptional repression of LacI leads to the synthesis of TreS-Mak activity and *per se* the synthesis of highly toxic maltose-1-phosphate that do not occur in BL21-AI strains. **(B)**. Recombinant his-tag TreS-Mak was purified on nickel affinity column and total protein of crude extract (CE), Flow through (FT), Washing steps (W1, W2, W3, W4) and Elution step (E1, E2, E3) were separated on 7.5% SDS-PAGE. Based in molecular weight standard, blue Coomassie staining revealed a polypeptide at 115 kD. **C** Superose 6 Increase 10-300 GL column (GE-Healthcare) pre-equilibrated with 140 mM NaCl, 10 mM orthophosphate pH 7.4 was calibrated with standard proteins (669; 440; 158; 43 and 13.7 kDa) and Blue Dextran. The determination of a partition coefficient (Kav) of 0.5 suggests an apparent molecular weight of 256 kD. **D**. Effect of imidazole concentration on maltokinase activity of TreS-Mak. Maltokinase activity incubated in a reaction buffer containing maltose (20 mM), ATP (20 mM) Mn^2+^ (10 mM) and various concentrations of imidazole at pH 8 (25 mM to 125 mM). The production of ADP is enzymatically determined after 40 minutes at 42°C (see material and methods). **E** and **F** pH and temperature optima of Mak activity were assayed with reaction buffer and same incubation time as described above. Mak activity was determined at pH 6, 7, 8 and 9 using imidazole (125 mM) as buffer at 30°C. Optimal temperature was inferred with reaction buffer containing 125 mM imidazole pH 7.

**S8 Figure:**
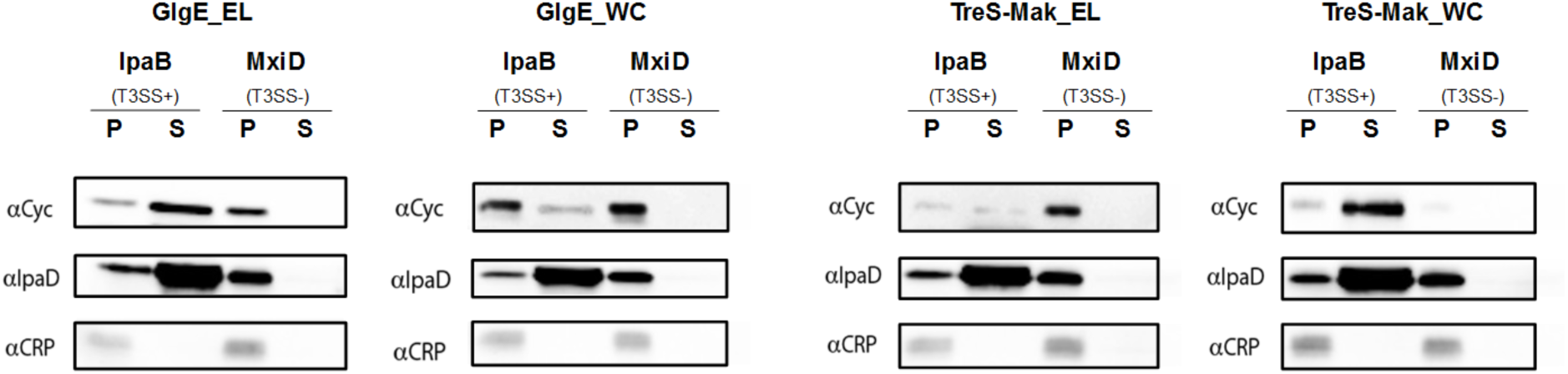
Heterologous secretion assay in *Shigella flexneri* of GlgE and TreS-Mak proteins. The first thirty amino acids at N-terminus extremity of each protein were fused with the reporter protein adenylate cyclase from *B. pertussis* (Cyc). Fused proteins are expressed in IpaB (T3SS+) and MxiD (T3SS-) strains of *S. flexneri* harboring a functional and a defective type-three secretion system, respectively. Western blot analyses were performed on both cell pellets (P) and supernatants (S) using adenylate cyclase antibodies (αCyc). In parallel, a secreted (IpaD) and a non-secreted protein (CRP) were selected as positive and negative controls, respectively. Both proteins were detected in cell pellets or supernatants using αCRP and αIpaD antibodies. Those preliminary results suggest that both GlgE and TreS-Mak proteins of *Estrella lausannensis* (EL) and *Waddlia chondrophila* (WC) are secreted by the type-three secretion system.

**S1 table:**
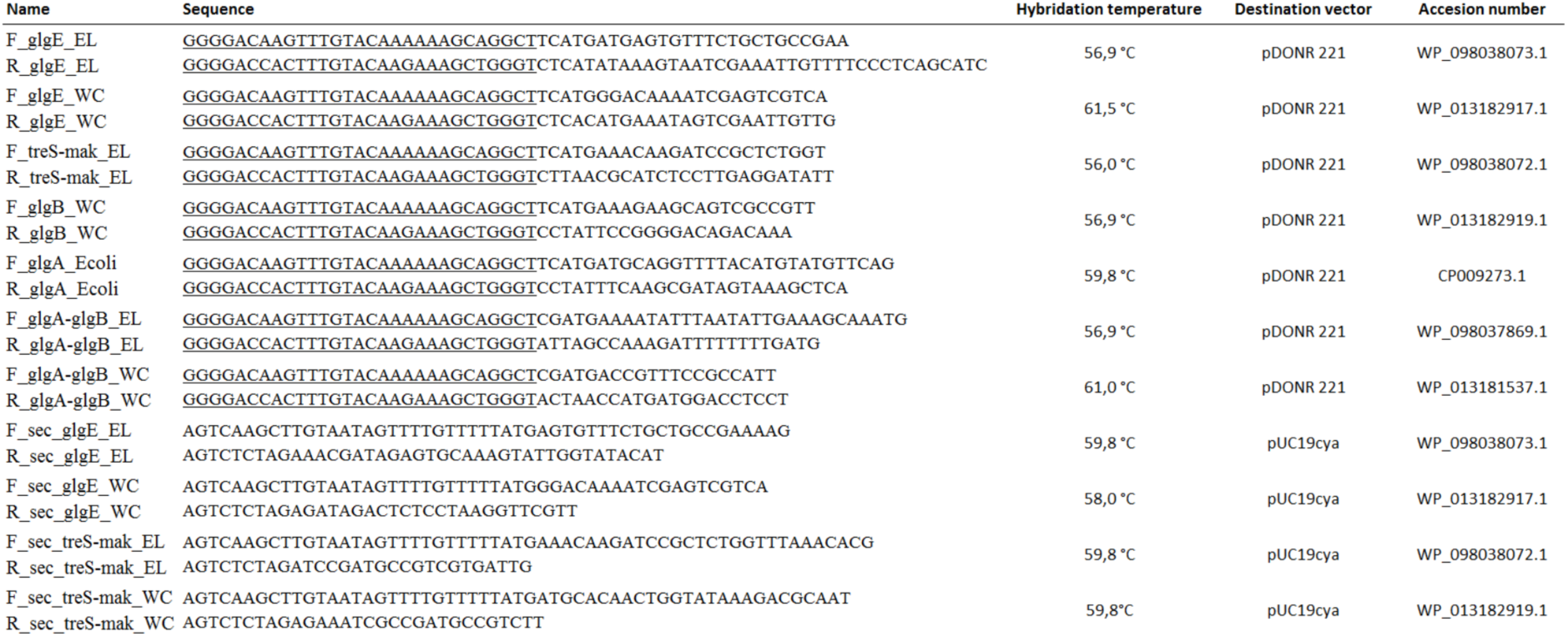
List of primers used for gene cloning involved in glycogen metabolism pathway of *E. lausannensis* and *W. chondrophila* and for heterologous secretion assay. Underlined nucleotides represent the attB sites added to the amplified genes that allow the cloning into pDONR 221 vectors following the recommendation of Thermofisher (Gateway^TM^).

**S2 Table:** Organization of *glg* genes of GlgC pathway across Chlamydial genomes. Genes encoding for ADP-glucose pyrophosphorylase (glgC); glycogen synthase (glgA), glycogen branching enzyme (glgB), glycogen phosphorylase (glgP) and glycogen debranching enzyme were listed according to their organization on chlamydial genomes. With a notable exception for glgC and glgP genes, which are often separated by one or two genes, most of glg genes are encoded more than 10 kpb from each other.

| Genome accession | species | organization of glg genes |  |  |  |  |
| --- | --- | --- | --- | --- | --- | --- |
| NZ_LT999999 | *Chlamydia suis* isolate 4-29b | X | P | C | A | B |
| CP000051 | *Chlamydia trachomatis* A/HAR-13 | X | P | C | A | B |
| AM884177 | *Chlamydia trachomatis* L2b/UCH-1/proctitis | A | B | X | P | C |
| CP021996 | *Chlamydia abortus* GN6 | C | B | X | P | A |
| CP001713 | *Chlamydia pneumoniae* LPCoLN | C | B | X | P | A |
| AE009440 | *Chlamydia pneumoniae* TW-183 | P | X | B | C | A |
| CR848038 | *Chlamydia abortus* S26/3 | C | B | X | P | A |
| AE015925 | *Chlamydia caviae* GPIC | C | B | X | P | A |
| CP002549 | *Chlamydia psittaci* 6BC | C | B | X | P | A |
| CP004033 | *Chlamydia pecorum* PV3056/3 | C | B | X | P | A |
| APJW01000000 | *Chlamydia ibidis* 10-1398/6 | C | B | X | P | A |
| ATNB01000000 | *Chlamydia ibidis* 10_1398_11 | X | P | A | C | B |
| MKSK01000000 | *Chlamydiales* bacterium 38-26 | A | B | X | C | P |
| CP019792 | *Chlamydia gallinacea* JX-1 | P | X | B | C | A |
| AE002160 | *Chlamydia muridarum* Nigg | A | B | X | P | C |
| AP006861 | *Chlamydia felis* Fe/C-56 Fe/C-56 | A | P | X | B | C |
| CP002608 | *Chlamydia pecorum* E58 | A | C | B | X | P |
| CP015840 | *Chlamydia gallinacea* 08-1274/3 | A | C | B | X | P |
| CP006571 | *Chlamydia avium* 10DC88 | B | X | P | A | C |
| FR872580 | *Parachlamydia acanthamoebae* UV-7 | C | P | X | B | A |
| ACZE01000000 | *Parachlamydia acanthamoebae* Hall's coccus | C | P | B | A | X |
| JSAM01000000 | *Parachlamydia acanthamoebae* OEW1 | C | P | X | B | A |
| AP017977 | *Neochlamydia* sp. S13 | A | X | B | P | C |
| JSAN01000000 | *Candidatus Protochlamydia amoebophila* EI2 | P | C | A | B | X |
| LN879502 | *Candidatus Protochlamydia naegleriophila* KNic | X | B | A | C | P |
| DNHV01000000 | *Parachlamydiales* bacterium | X | B | A | C | P |
| NZ_FCNU00000000 | *Parachlamydia* sp. C2 | B | X | A | C | P |
| CAAJGQ000000000 | *Rhabdochlamydia* sp. T3358 | X | P | A | C | B |
| NZ_CCJF000000000 | *Protochlamydia massiliensis* | A | C | P | X | B |
| NZ_BAWW000000000 | *Parachlamydia acanthamoebae* Bn9 | P | B | A | C | X |
| FR872582 | *Simkania negevensis* Z main | C | P | B | X | A |

